# On demand MyD88 oligomerization is controlled by IRAK4 during Myddosome signaling

**DOI:** 10.1101/2020.09.03.280917

**Authors:** Rafael Deliz-Aguirre, Fakun Cao, Fenja H. U. Gerpott, Nichanok Auevechanichkul, Mariam Chupanova, YeVin Mun, Elke Ziska, Marcus J. Taylor

## Abstract

A recurring feature of innate immune receptor signaling is the self-assembly of signaling proteins into oligomeric complexes. The Myddosome is an oligomeric complex that is required to transmit inflammatory signals from TLR/IL1Rs and consists of MyD88 and IRAK family kinases. However, the molecular basis for how Myddosome proteins self-assemble and regulate intracellular signaling remains poorly understood. Here, we developed a novel assay to analyze the spatiotemporal dynamics of IL1R and Myddosome signaling in live cells. We found that MyD88 oligomerization is inducible and initially reversible. Moreover, the formation of larger, stable oligomers consisting of more than 4 MyD88s triggers the sequential recruitment of IRAK4 and IRAK1. Notably, genetic knockout of IRAK4 enhanced MyD88 oligomerization, indicating that IRAK4 controls MyD88 oligomer size and growth. MyD88 oligomer size thus functions as a physical threshold to trigger downstream signaling. These results provide a mechanistic basis for how protein oligomerization might function in cell signaling pathways.

## Introduction

The innate immune system is a form of host defense that rapidly responds to infection and disease (Medzhitov and Janeway, 2000). Central to an innate immune response are diverse receptor families that are germline encoded and recognize the molecular signals of infection and disease (Akira et al., 2006). A unifying property of innate immune receptor signaling pathways is the self-assembly of signaling proteins into large macromolecular complexes (Kagan et al., 2014). Structural and molecular characterization revealed that these macromolecular assemblies are oligomeric, with signaling effectors able to polymerize into structurally defined complexes (Wu, 2013), and are collectively referred to as supramolecular organizing centers (SMOCs) (Kagan et al., 2014).

Unlike receptor Tyrosine kinases or G-protein coupled receptors, many innate immune receptors are not enzymatically active nor directly linked to secondary messengers, such as calcium or cAMP (Wu, 2013). Furthermore, the signaling effectors that bind to activated receptors and self-assemble into SMOCs do not contain enzymatic activity. Therefore, innate immune receptors such as inflammasome receptors, tumor necrosis factor (TNF) receptors, Toll-like receptors (TLR), and Interleukin 1 receptors (IL1R) cannot simply transduce signals by upregulating enzymatic activity (Kagan et al., 2014; Wu, 2013). Thus, a model for SMOC signalling is that these macromolecular complexes are inducible platforms that form on demand and activate signalling by concentrating and activating downstream enzymatic effectors (Tan and Kagan, 2019). However, how is receptor triggered oligomerization controlled to accurately and rapidly transduce a signal? How large must a SMOC oligomer be and how long must it persist to achieve downstream signaling?

One such SMOC, the Myddosome, is a macromolecular complex consisting of helical oligomers of MyD88, and kinases of the IL1 receptor-associated kinase (IRAK) family (Lin et al., 2010; Motshwene et al., 2009). The Myddosome mediates signaling from the TLR/IL1R superfamily (Gay et al., 2014). Members of the TLR/IL1R superfamily are critical mediators of a protective innate immune response and are characterized by the presence of a cytoplasmic Toll-interleukin1 receptor (TIR) domain (O’Neill, 2008). IL1Rs respond to inflammatory cytokines of the IL1 family (Dinarello, 2009), whereas TLRs respond to microbial- and viral-associated molecules (Gay et al., 2014). The Myddosome biochemically interacts with activated TLRs/IL1Rs via the TIR domain containing cytoplasmic adapter MyD88 (Adachi et al., 1998; O’Neill and Bowie, 2007). MyD88 has no intrinsic enzymatic activity and contains an N-terminal Death Domain (DD) and a C-terminal TIR domain (Hardiman et al., 1996). MyD88 biochemically interacts with TLRs/ILR1s via heterotypic TIR domain interactions (Medzhitov et al., 1998), and self-assembles via its DD into helical oligomers (Lin et al., 2010). It is these helical MyD88 oligomers which couple to enzymatic activity by co-assembling with the DD-containing Ser/Thr kinases of the IRAK family.

Myddosome assembly is thought to be triggered by TLR/ILR1 ligand binding and receptor dimerization. This is believed to stimulate the recruitment and assembly of Myddosome components at the plasma membrane. Consistent with this model, assembled Myddosomes containing MyD88 and IRAK kinases can only be isolated by biochemical pull-downs from LPS-activated macrophages (Bonham et al., 2014). Structural studies on purified Myddosomes revealed a hierarchical order of stacked death domain oligomers consisting of 6 MyD88s, followed by 4 IRAK4s, and 4 IRAK2s (Lin et al., 2010). This organization suggests a sequential order of assembly, i.e. MyD88 polymerization triggers the recruitment of IRAK4 followed by IRAK2 (or the functionally redundant IRAK1 (Kawagoe et al., 2008)). However, purified MyD88 death domains and full length protein can polymerize into helical open-ended filaments *in vitro* (Moncrieffe et al., 2020; Motshwene et al., 2009; O’Carroll et al., 2018). It remains unclear how the size of MyD88 oligomers is controlled *in vivo.* In conclusion, while the major components of Myddosome signaling have been identified, it is currently not clear how the co-assembly of MyD88 and IRAK kinases is controlled at a precise time and place within live cells.

Visualizing the spatiotemporal dynamics of Myddosome assembly in living cells could unveil hitherto hidden mechanisms of TLR/IL1R signal transduction. Live cell analysis of Myddosome dynamics has been limited to individual proteins such as MyD88 or IRAK1 (Latty et al., 2018; Vayttaden et al., 2019). This has made it difficult to determine how the multiple proteins required for TLR/IL1R signaling are temporally coordinated and to identify precise stages in Myddosome assembly. Here, we develop a new live imaging approach to directly visualize Myddosome formation in response to IL1β stimulation in EL4 cells. We engineer precise fluorescent protein fusions of Myddosome proteins at endogenous gene loci using CRISPR/Cas9. By simultaneously imaging and quantifying multiple signaling reactions, we discovered that the formation of larger MyD88 oligomers functions as a signaling threshold to trigger IRAK kinase recruitment to the cell surface and IRAK4 regulates MyD88 oligomerization. Collectively, these results highlight how protein oligomerization can transduce biochemical signals. This provides a conceptual framework for understanding SMOC assembly in diverse innate immune receptor signaling pathways.

## Results

### Membrane-tethered IL1β triggers the relocalization of MyD88 to the cell surface and nuclear translocation of RelA

To image the molecular dynamics of Myddosome signaling, we used CRISPR/Cas9 gene editing to generate monoclonal cell lines with MyD88 tagged at the endogenous gene locus with a C-terminus monomeric enhanced green fluorescent protein (GFP) (Fig. S1A-D). We performed gene editing in the mouse lymphoma T cell line EL4.NOB1 (referred to as EL4 cells, see Material and Methods). We selected EL4 cells as an experimental system because they are highly responsive to IL1 and have previously been used to study IL1 signaling (Bird et al., 1988; O’Neill et al., 1990). Our knockin strategy enabled us to limit overexpression artifacts and quantitatively compare cells and measurements across experiments.

To stimulate IL1R signaling and Myddosome formation in live EL4 cells, we developed planar supported lipid bilayers (SLBs) functionalized with freely diffusing IL1β (Fig. 1A). Since IL1R signals in response to soluble and membrane-bound isoforms of IL1(Kaplanski et al., 1994), we reasoned that IL1 tethered to SLBs reconstituted the IL1R signaling at cell-cell contact sites. Finally, the planar geometry of SLBs can be combined with Total Internal Reflection (TIRF) microscopy to directly visualize signaling reactions at the cell surface (Biswas and Groves, 2019).

**Figure 1.**
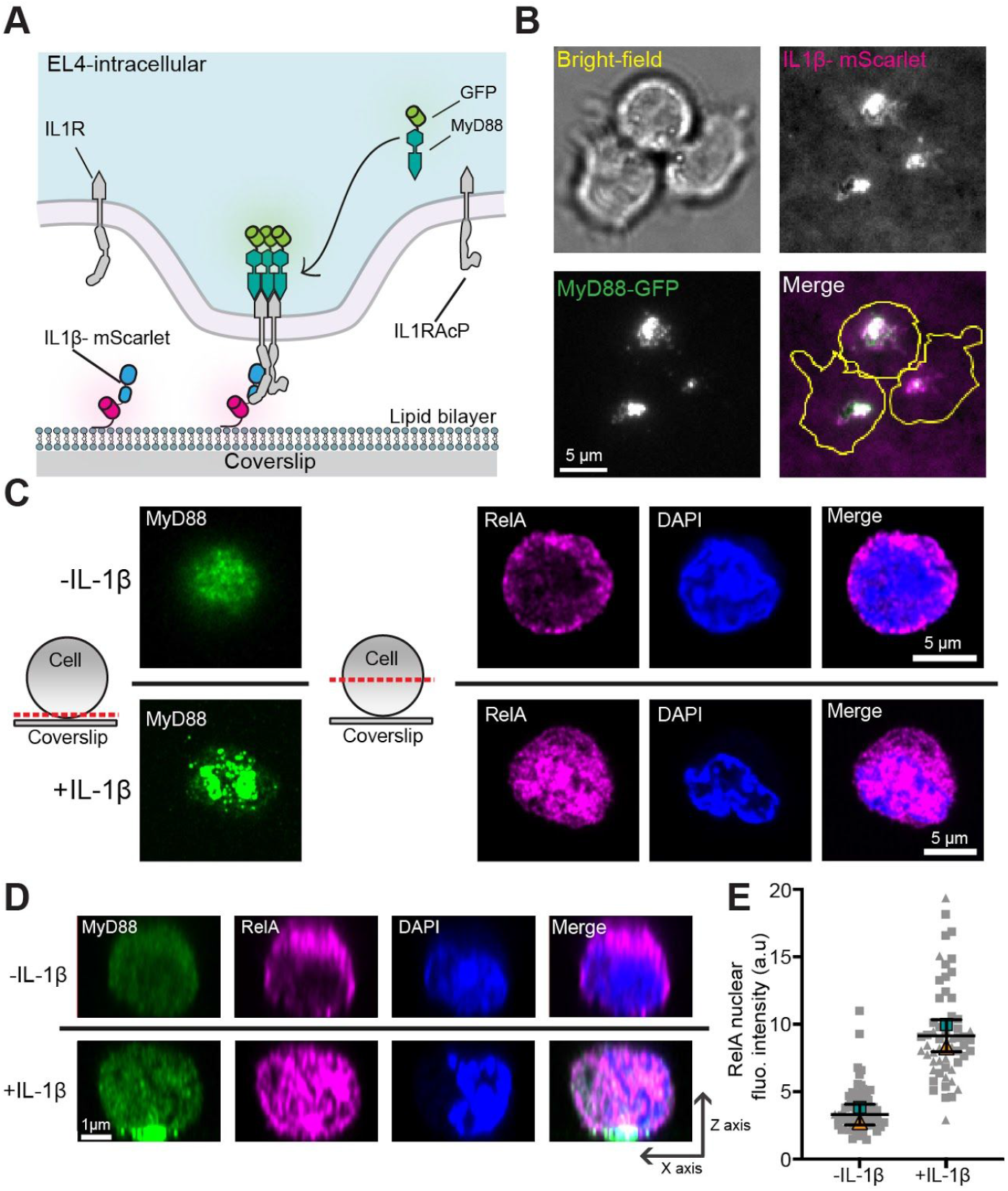
Membrane-tethered IL1β triggers the relocalization of MyD88 to the cell surface and nuclear translocation of RelA. **A) The schematic of a supported lipid bilayer (SLB) functionalized with IL1β labelled with mScarlet.** **B) TIRF and brightfield microscopy images of EL4 cells expressing MyD88-GFP after landing on a SLB functionalized with IL1β-mScarlet.** Clusters of IL1β-mScarlet formed at the cell-SLB interface. MyD88-GFP was recruited to clusters of IL1β-mScarlet. Scale bar, 5 μm. **C-D) IL1β tethered to SLBs activates RelA nuclear translocation in EL4 cells.** To measure activation of NF-kB, EL4 cells were fixed (45 mins after SLB contact) and stained for MyD88-GFP (green); RelA (magenta); DAPI staining of nuclei (blue). Cells were imaged with confocal microscopy. (C) Schematic shows the position of the confocal micrograph slice. Cells in contact with SLB functionalized with IL1β relocalise RelA to the nucleus. Scale bar, 5 μm. (D) Reconstructed axial view of cells shown in (C), showing the localization of MyD88 to the cell-SLB contact zone and RelA to cell nucleus under IL1β stimulation. Scale bar, 1 μm. **E) Quantification of RelA nuclear staining intensity**. Bar represents mean ± SEM from n = 2 experimental replicates. Scatter plot symbols representing independent replicates, smaller gray symbols represent single-cell measurements and superimposed larger symbols represent the averages from experimental replicates.

We pipetted EL4-MyD88-GFP cells into chambers containing IL1β-mScarlet labeled SLBs (Fig. 1A). Cells were allowed to settle for 20 mins before being imaged using TIRF and bright-field microscopy. We observed that IL1β-mScarlet was clustered at the EL4 cell-bilayer interface. MyD88-GFP was recruited to this interface and assembled into large fluorescent clusters. These macromolecular clusters co-localized with the IL1β-mScarlet clusters (Fig. 1B).

We determined whether membrane-tethered IL1β could activate downstream signaling outputs. Activation of IL1R triggers the stimulation of NF-kB protein complex and the translocation of the p65-RelA subunit into the nucleus. We analyzed the subcellular distribution of RelA in EL4-MyD88-GFP cells incubated with IL1β-labeled supported membranes for 60 mins before being fixed and stained with anti-RelA. The cellular volume was then imaged with confocal microscopy (Fig. 1C-D). Cells incubated with IL1β-labeled SLBs clustered MyD88-GFP at the cell-bilayer interface. The accumulation of MyD88-GFP was associated with the localization of RelA to the nucleus (Fig. 1C-D).

In EL4 cells incubated with unlabeled supported membranes, MyD88-GFP remained diffuse throughout the cytoplasm and RelA was excluded from the nucleus, thereby resulting in lower nuclear staining intensities (Fig. 1E). Similar to NF-kB activation, we found that EL4 cells incubated with IL1β-labeled SLBs had increased levels of phospho-p38 (Fig. S1E-G). Thus, IL1β tethered to supported membranes can activate NF-kB and MAPK p38 signaling as well as the re-localization of MyD88 to the cell surface.

### MyD88-GFP puncta form at the cell surface and colocalize with clusters of IL1R-bound IL1β

We examined the temporal dynamics of MyD88-GFP in EL4 cells stimulated with IL1β. EL4-MyD88-GFP cells were applied to SLBs and imaged using TIRF microscopy (Fig. 2A). We observed that these puncta were mobile at the cell-bilayer interface, and after several mins, coalesced into larger clusters (Fig. 2A). MyD88-GFP puncta were dynamic and underwent both fusion and fission (Fig. S2). We quantified the formation of MyD88-GFP puncta as a function of time after cell landing. Within five mins of contacting the IL1β-functionalized supported membrane, EL4 cells rapidly formed many (5-25) MyD88-GFP puncta (Fig. 2B).

**Figure 2.**
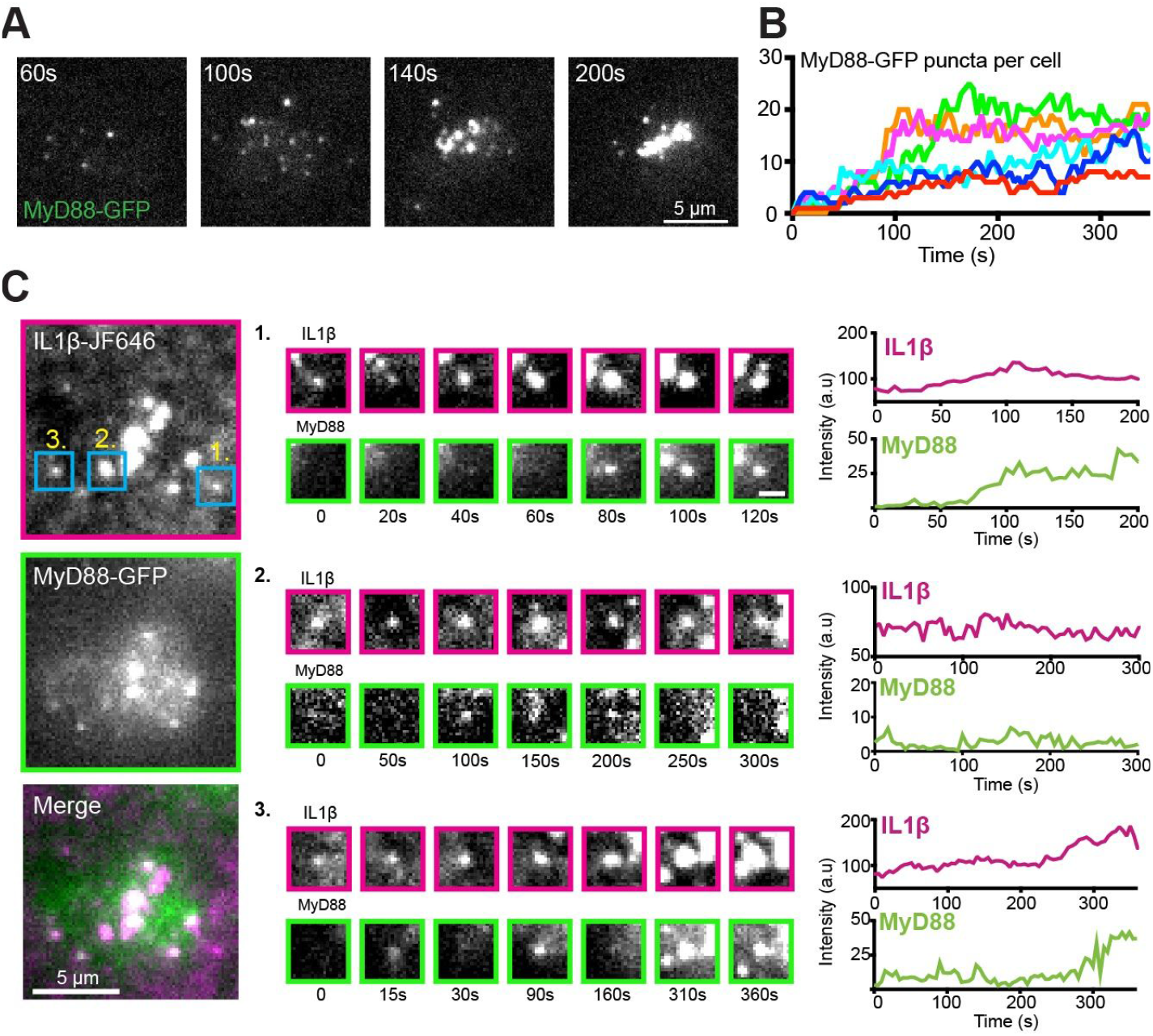
MyD88-GFP puncta form at the cell surface and colocalize with clusters of receptor-bound IL1β. **A) Time-lapse TIRF images of MyD88-GFP showing the formation of MyD88-GFP puncta over time at the cell surface**. MyD88-GFP puncta coalescence into larger clusters at the cell-SLB contact interface. Scale bar, 5 μm. **B) Formation over time of MyD88-GFP puncta for individual cells (colors)**. t = 0 is defined as the point of image acquisition, generally within 1–2 min of adding cells to the SLB. **C) TIRF images of IL1β-JF646 and MyD88-GFP showing Myddosome formation at clusters of IL1β-bound IL1R**. Scale bar, 5 μm. Regions of interest (blue boxes) show three IL1β-receptor clusters (labeled 1–3). Region of interest (ROI) scale bar, 1.5 μm. Fluorescence-intensity time series from the IL1β-JF646 and MyD88-GFP of the three IL1β clusters are shown on the right.

We investigated the recruitment of MyD88 to IL1β-IL1R complexes. To observe IL1β-IL1R engagement, we engineered a Halo-tag version of IL1β and labeled it with the photostable organic fluorophore JF649. Using two-color TIRF microscopy, we imaged both cellular MyD88-GFP and IL1β-JF649 on the SLBs. When EL4 cells contacted the supported membrane, we observed the homogeneous distribution of IL1β-JF646 reorganized into microclusters (Fig. 2C and Movie S1). Many IL1β microclusters increased in fluorescence intensity over time, indicating the addition of newly IL1R-bound IL1β-JF649. IL1β microclusters first appeared as diffraction-limited clusters, but over time coalesced into larger non-diffraction patch-like structures. MyD88 spots formed at clusters of IL1β at the cell-bilayer interface. The formation of IL1β clusters preceded the recruitment and assembly of MyD88 into puncta (Fig. 2C). From these data, we conclude that IL1β binding to IL1R stimulates the recruitment of MyD88 to the plasma membrane. At the plasma membrane, MyD88 then assembles into puncta that colocalize with clusters of IL1β-bound IL1R.

We observed that MyD88 had heterogeneous recruitment dynamics to clusters of IL1β-bound IL1R. We observed the formation of stable MyD88 puncta (“stable” defined as persisting for more than one min, Fig. 2C example 1). The stable MyD88 puncta appeared ~1-2 mins after the formation of an IL1β cluster. We also observed the formation of transient MyD88 foci that formed at the cell surface. These transient foci were generally dimmer than the stable MyD88 puncta and did not increase in fluorescence intensity (Fig. 2C, example 2). In some cases, we observed both types of dynamics at the same clusters of IL1R-bound IL1β. In these instances, the transient MyD88-GFP foci preceded the formation of a stable MyD88 punctum (Fig. 2C, example 3).

### MyD88-GFP forms both transient and stable macromolecular assemblies

We hypothesized that these heterogeneous recruitment dynamics reflected the process of MyD88 self-assembly into oligomers. To understand this process in greater detail, we used automated particle tracking to obtain an unbiased dataset of MyD88 assemblies and then analyzed their lifetimes and intensities (Movie S2). To estimate the copy number of MyD88 within the MyD88-GFP puncta, we calibrated our TIRF setup using purified GFP. Congruent with our previous observation (Fig. 2C), we observed two classes of MyD88-GFP puncta at the cell surface with distinct lifetimes and intensity traces. The first class contained the transient foci that showed a minimal increase in fluorescent intensity. The second class contained foci that increased in fluorescent intensity becoming bright stable MyD88-GFP puncta (Fig 3A, see example of each dynamic, and Movie S2). We observed that MyD88-GFP foci corresponding to the first class had fluorescent intensities corresponding to 1-3x the mean intensity of GFP. MyD88 puncta that belonged to the second class had an initial fluorescence intensity equivalent to 1-3x GFP. In contrast to the shorter-lived MyD88 assemblies, these structures increased to a fluorescent intensity in a manner consistent with >6x GFP mean intensity (Fig. 3A, dashed lines on the intensity time traces).

**Figure 3.**
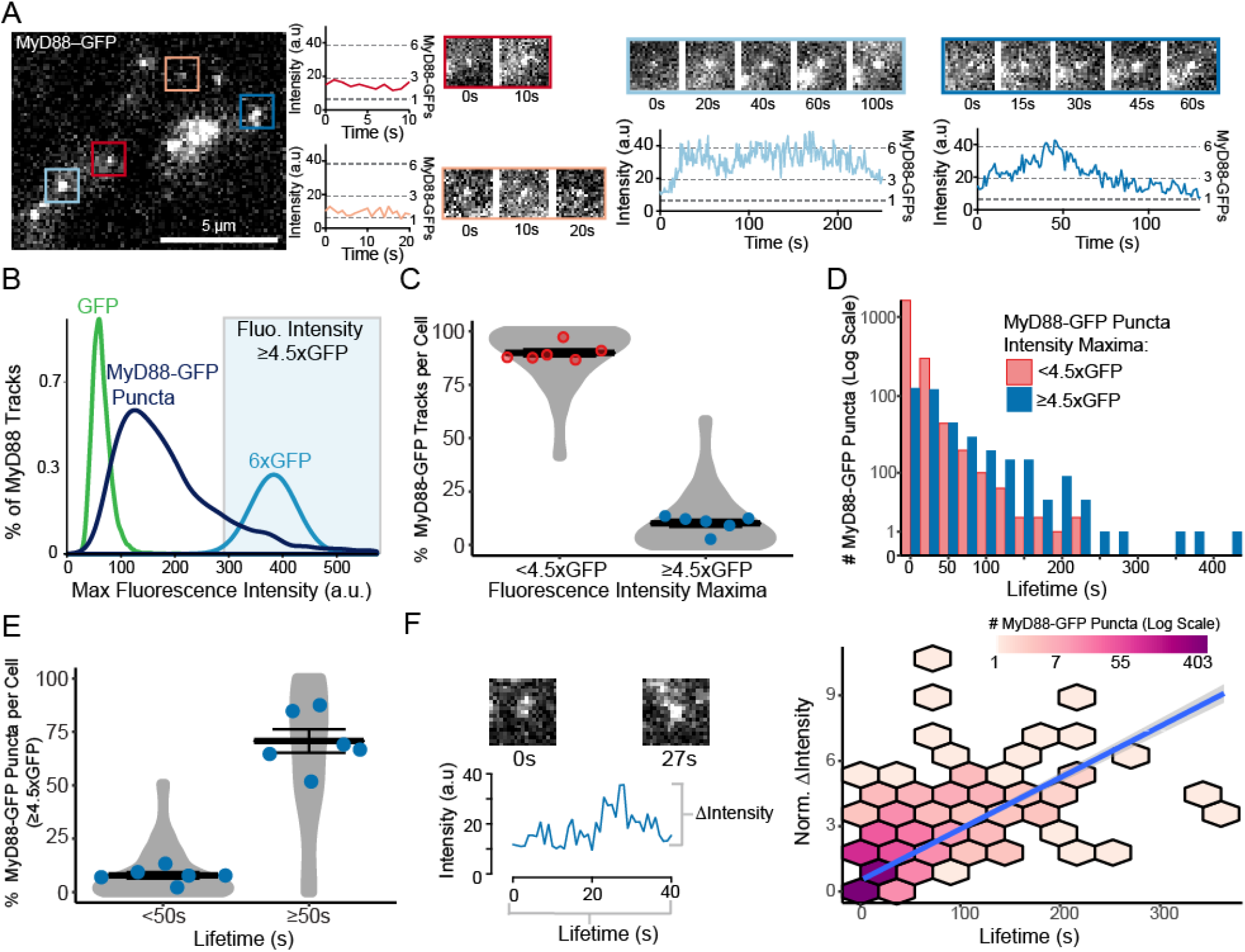
Analysis of MyD88-GFP puncta dynamics, lifetime and size. **A) TIRF images of MyD88-GFP in an EL4 landing on an IL1β functionalized SLB.** Overlaid coloured boxes highlight examples of individual Myddosomes. Scale bar, 5μm. Red and orange ROI show the fluorescence intensity time series (left panel) and TIRF images (right panel) from myddosomes that are short lived (<50 s) and dim (<3x GFP average intensity). Blue ROIs show fluorescence intensity time series (bottom panel) and TIRF images (top panel) from two example MyD88-GFP puncta that grow in fluorescence intensity and have long life time (≥50 s). The dashed gray lines on intensity plots mark the quantal MyD88-GFP fluorescence intensities, estimated from single GFP fluorophores. **B) MyD88-GFP puncta have a wide distribution of sizes**. Density plot of the maximum fluorescent intensity of MyD88-GFP puncta (dark blue, n = 2,422 tracked MyD88-GFP particles from 14 cells) compare to single molecules of GFP (green, n = 397 GFP particles) and estimated intensity distribution of a 6x GFP multimer (light blue). A 6x GFP multimer was estimated by fitting the 1x GFP intensity distribution to a Gaussian (see Methods). Shaded blue region designates fluorescent intensity values greater than 4.5x GFP. **C) Quantification of the proportion (%) of MyD88-GFP puncta per cell that have a maximum fluorescence intensity <4.5xGFP or ≥4.5xGFP**. Violin plots show the distribution of the cell data. Data points superimposed on violin plots are the averages of independent experiments. Bars represent mean ± SEM (n = 6 experimental replicates, with 6-24 cells measured per replicate). **D) Distribution of MyD88-GFP puncta dwell time**. Myddosomes were classified by maximum fluorescence intensity< or ≥ 4.5x GFP (n = 2,037 <4.5x GFP versus n = 385 ≥4.5x GFP tracked MyD88-GFP puncta combined from 14 cells). Myddosomes with greater dwell times tend to have a maximal fluorescent intensity that is ≥4.5x GFP. **E) Quantification of the proportion (%) per cell of MyD88-GFP puncta with an intensity maxima ≥ 4.5x GFP categorized by lifetimes < or ≥ that 50 s**. Violin plots show the distribution of the cell data. Data points superimposed on the violin plots are the averages from independent experiments. Bars represent mean ± SEM (n = 6 experimental replicates, with 6-24 cells measured per replicate). **F) Correlation between growth in intensity and lifetime of MyD88-GFP puncta.** Left panel: TIRF images and intensity trace of a representative MyD88-GFP puncta. The change in intensity for each tracked MyD88-GFP puncta was calculated as maximum intensity subtracted by the initial intensity and normalized to the intensity of GFP. Right panel: 2D histogram of MyD88-GFP puncta lifetime versus changed in fluorescent intensity. MyD88-GFP puncta that increase in fluorescent intensity tend to have progressively longer dwell times. Linear fit is shown as a blue line with the 95% confidence interval shown in gray (Spearman’s rank correlation coefficient R = 0.59, p<.001, n = 1,763 MyD88 tracks).

We systematically examined the distribution of MyD88 oligomer sizes. We plotted the distribution of maximum intensities of tracked MyD88 particles and compared this distribution to the fluorescent intensity of single GFP fluorophores. Given the Myddosome crystal structure contains 6x MyD88s (Lin et al., 2010), we also estimated the fluorescent intensity distribution for a particle containing 6xGFP molecules (Fig. 3B, see Methods). The MyD88-GFP puncta distribution suggested a broad size distribution of detected MyD88 assemblies (Fig. 3B, 2422 puncta, collated from 14 cells, also see Fig. S3A for additional replicates). A minority of MyD88 puncta had a fluorescence intensity equivalent to 6x multimers. In contrast, we find the majority of MyD88 puncta consisted of ~2-3 MyD88 monomers.

We devised a quantitative classification of MyD88 puncta size (Fig 3B). Based on the distribution of single GFP fluorophores, we used a maximum intensity threshold of 4.5xGFP to classify puncta as large MyD88 oligomers (e.g. greater than 4 MyD88s). We subsequently analyzed the proportion of MyD88 puncta with a max intensity of < or ≥ 4.5xGFP, i.e. defined as small or large MyD88 assemblies, respectively. We applied this classification metric to all particle-tracked MyD88-GFP puncta detected within single EL4 cells (see Movie S2). We found that on average, less than 12% of MyD88 puncta had a maximum intensity of ≥ 4.5x (Fig. 3C and Fig. S3B). The majority of MyD88 puncta had intensities of <4.5xGFP (90 ± 10% of MyD88-GFP puncta per cell, mean ± SEM, calculated from six independent replicates, Fig. 3C and Fig. S3B). Lifetime analysis revealed that many of these small MyD88 oligomers had short dwell times of between 3-10 s (Fig. 3D and Fig S3C). In contrast, larger MyD88 oligomers had longer lifetimes and could persist for more than 100 s (Fig. 3D).

We classified MyD88 puncta as short or long-lived (defined as < or ≥50 s) and analyzed the proportion of large assemblies per cell in each lifetime category. We found that 8% of MyD88 puncta per cell with lifetimes <50 s were large assemblies. In contrast, 68% of MyD88 puncta per cell with lifetimes ≥50 s were large oligomers (Fig. 3E, and Fig. S3D). Thus, long-lived and stable MyD88 puncta tend to be larger assemblies.

We hypothesized that if MyD88 oligomerization is inducible, the stable larger MyD88 puncta would start as small assemblies and then grow in intensity during oligomerization. This hypothesis is consistent with a nucleation and growth process and would suggest there is a correlation between growth in intensity and lifetime. Small unstable MyD88 oligomers that fail to grow would rapidly disassemble, while oligomers that grow beyond a certain size would persist for longer at the cell surface. Consistent with this model, we found that MyD88-GFP puncta that increased in fluorescent intensity (defined as maximum intensity subtracted by the initial intensity) correlated with a longer lifetime at the cells surface (Spearman’s rank correlation coefficient, R = 0.59, calculated from 1,763 MyD88 puncta combined from 14 cells, Fig. 3F and Fig S3E for experimental replicates).

Thus, we find that MyD88-GFP puncta are oligomers of MyD88 nucleating in response to IL1R activation with the majority of these oligomers being transient and small (e.g. consisting of 2-3 MyD88 monomers, Fig. 3B). However, a portion of the nucleated oligomers recruit additional MyD88 and grow to become larger stable oligomers that can persist at the cell surface for lifetimes of ≥50 s.

### IRAK4 and IRAK1 are recruited to larger and kinetically stable MyD88 oligomers

We asked whether the size of the MyD88 oligomer regulated the recruitment of downstream effectors IRAK4 and IRAK1 to the cell surface. Because microscopy studies only imaged MyD88 within the TLR/IL1R signaling pathway (Latty et al., 2018; Latz et al., 2002), how MyD88, IRAK4, and IRAK1 cooperate within the TLR/IL1R signaling pathway remains unknown. To overcome this limitation, we used CRISPR/Cas9 to engineer monoclonal EL4 cell lines to express both MyD88-GFP and either IRAK4 or IRAK1 fused to the red fluorescent protein mScarlet-I in their endogenous genetic loci (Fig. S4A-C). We first analyzed the temporal dynamics of MyD88 and IRAK4 using multi-color TIRF microscopy in EL4 cells stimulated with our IL1β-functionalized SLB system. IRAK4-mScarlet, like MyD88, had a punctate localization pattern at the cell surface and also colocalized with MyD88 (Fig. 4A, and Movie S3). However, we observed that only a subset of MyD88 spots localized with IRAK4 at the plasma membrane (see Fig. 4A and Movie S3). Our analysis of the temporal dynamics uncovered that MyD88 puncta appeared at the cell surface before the recruitment of IRAK4 (an example time series is given in Fig. 4A).

**Figure 4.**
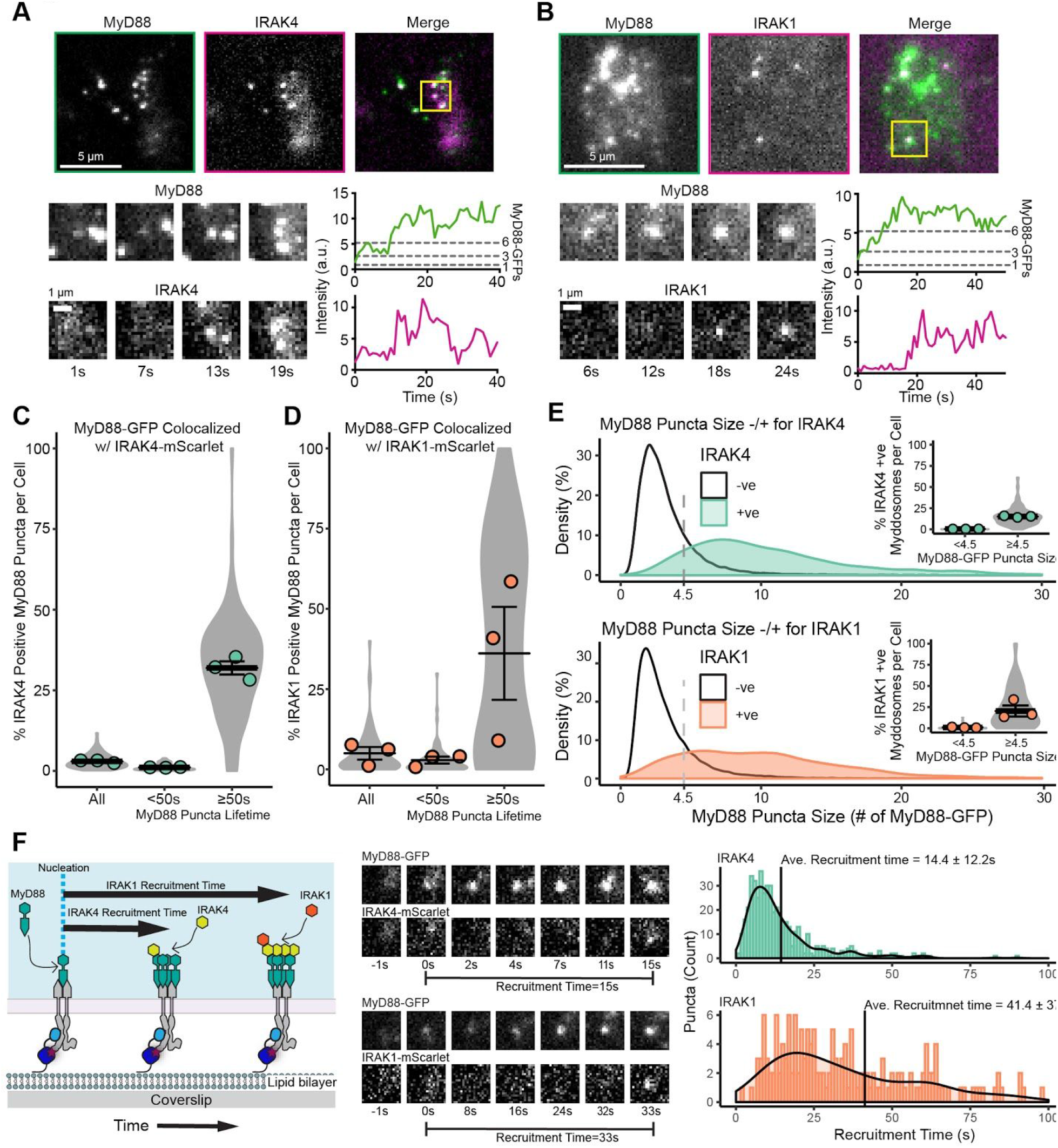
IRAK4 and IRAK1 are recruited to larger MyD88 oligomers. **A-B) Endogenously expressed IRAK4-mScarlet and IRAK1-mScarlet is recruited to MyD88-GFP puncta.** Top panel, TIRF images of monoclonal endogenously expressed MyD88-GFP and IRAK4-mScarlet (A) or IRAK1-mScarlet (B) EL4 cell lines. Scale bar, 5 μm. ROIs (yellow box, merge image) shows an example of a MyD88-GFP spot colocalized with IRAK4-mScarlet (A) or IRAK1-mScarlet. Bottom panels, time-series TIRF images from the ROI (left panel) and fluorescence-intensity time series (right panel) of MyD88-GFP and IRAK4-mScarlet (A) or IRAK1-mScarlet (B). Scale bar, 1 μm. **C-D) Longer-lived MyD88 punta have a greater colocalization with IRAK4 and IRAK1.** Quantification of the % of MyD88-GFP puncta per cell that colocalize with IRAK4 (C) or IRAK1 (D) for all puncta and puncta with lifetimes <50 or ≥50 s. Violin plots show the distribution of individual cell measurements. Colored dots superimposed on violin plots correspond to the average value in the independent experiments (n = 3 for IRAK4 and IRAK1, each replicate encompasses measurements from 16-34 cells). Bars represent mean ± SEM. **E) Larger oligomers of MyD88 colocalize with IRAK4 and IRAK1.** Density plot showing the distribution of MyD88 oligomer size (number of MyD88-GFP monomers is derived from the–maximum intensity divided by the average intensity of GFP) for MyD88 puncta that are positive (+ve) or negative (-ve) for IRAK4 (top) or IRAK1 (bottom). Inset: quantification of the % of MyD88-GFP puncta per cell that colocalize with IRAK4 or IRAK1 with a maximum intensity of < 4.5x GFP or ≥ 4.5x GFP. Violin plots show the distribution of individual cell measurements. Colored dots superimposed on the violin plots correspond to the mean value for the independent experiments (n = 3, IRAK4; n = 3, IRAK1). Bars represent mean ± SEM. **F) Analysis of the recruitment time shows the sequential recruitment of IRAK4 and IRAK1 during Myddosome assembly**. Recruitment time was defined as the time interval from Myddosome nucleation (e.g. t = 0 s when MyD88-GFP puncta appears) to the appearance of IRAK4/1-mScarlet at the cell surface. Middle panel, time series of TIRF images showing MyD88-GFP nucleation followed by IRAK4-mScarlet (top time series, IRAK4 initiation time = 15 s) and IRAK1-mScarlet (bottom time series, IRAK1 initiation time = 33 s) recruitment. Histogram of IRAK4 (n = 482 recruitment events, combined from 30 cells) and IRAK1 (n = 170 recruitment events, combined from 40 cells) initiation times overlayed with the density plot of the distribution. Black horizontal lines on the histograms denote the average initiation time for IRAK4 and IRAK1 (mean ± SD).

We then analyzed the temporal dynamics of MyD88 and IRAK1. Reminiscent of the IRAK4 localization puncta, IRAK1 had a punctate pattern at the cell surface. Only a subset of MyD88 spots localized with IRAK1 (see Fig 4B and Movie S4). Time series analysis revealed that IRAK1 puncta only appeared at the cell surface after the formation of the MyD88 puncta (Fig. 4B). From these data, we concluded that MyD88 is recruited before IRAK4/1 and only a subset of MyD88 assemblies recruit IRAK4/1.

To determine which properties of MyD88 assemblies trigger IRAK4 recruitment, we quantified the percentage of MyD88-GFP puncta that colocalized with a puncta of IRAK4-mScarlet. We found that 3% (Fig. 4C, n = 3 experimental replicates, >30 cells per replicate) of MyD88 puncta per cell colocalized with IRAK4. To assess whether longer-lived MyD88 assemblies were more likely to recruit IRAK4, we compared the IRAK4 recruitment to MyD88 puncta with lifetimes of < or ≥ than 50 s. Only 1.0% of MyD88 puncta per cell with a lifetime of <50 s recruited IRAK4-mScarlet. In stark contrast, 32% of MyD88 puncta per cell with a lifetime of ≥50 s recruited IRAK4-mScarlet (Fig. 4C, n = 3 independent replicates).

We repeated this analysis using IRAK1-mScarlet and determined that 5.0% of MyD88-GFP puncta colocalized with a puncta of IRAK1-mScarlet, but 36% of MyD88-GFP puncta with a lifetime of ≥50 s colocalized with IRAK1-mScarlet (Fig. 4D, n = 3 experimental replicates). Interestingly, only 3.0% of MyD88-GFP puncta with a lifetime of <50 s colocalized with IRAK1-mScarlet. Thus the long-lived MyD88-GFP assemblies more efficiently recruit IRAK4 and IRAK1 to the cell surface.

We also hypothesized that the size of a MyD88 oligomer regulates its association with downstream signaling effectors. To test this hypothesis, we analyzed the maximum intensity of MyD88-GFP particles that colocalized with IRAK4-mScarlet as compared to those that did not colocalize. We found that IRAK4-positive MyD88 particles were brighter than IRAK4-negative MyD88 particles (Fig. 4E). When we normalized the MyD88 intensity to GFP, we estimated that IRAK4 was recruited to larger MyD88 oligomers that had an average size of 11.0x MyD88-GFP. In comparison, MyD88 particles negative for IRAK4 had an average intensity of 3.2x MyD88-GFP (Fig. S5B). We repeated this analysis with IRAK1 and found that IRAK1-mScarlet positive MyD88 puncta were brighter than IRAK1-mScarlet negative MyD88 particles (Fig. 4E). IRAK1-positive MyD88-GFP puncta had an average size of 12x MyD88-GFP, whereas negative-MyD88 puncta had an average size of 3.7x MyD88-GFP (Fig. S5D). Consistent with this analysis, a greater proportion of larger MyD88 oligomers colocalized with IRAK4 and IRAK1 (inset Fig. 4E). Thus, IRAK4 and IRAK1 are recruited to larger, stable assemblies of MyD88.

### The Myddosome forms by the sequential recruitment of IRAK4 and IRAK1

Structural studies have revealed that within the Myddosome core complex, MyD88 interacts directly with IRAK4, which in turn interacts with IRAK1 (Lin et al., 2010). These studies predict that Myddosomes assemble sequentially. However, this model of assembly has never been visualised in live cells. The experiments above found that stable larger MyD88-GFP oligomers recruited both IRAK4 and IRAK1 (Fig. 4C-E). This led us to directly measure the recruitment kinetics of IRAK4 and IRAK1 to individual MyD88-GFP oligomers.

To analyze the kinetics of formation, we measured the time from MyD88 nucleation to the appearance of IRAK4 or IRAK1 (schematic, Fig. 4F). We defined this time measurement as the “recruitment” time. This analysis defined distinct recruitment kinetics for IRAK4 and IRAK1. Both IRAK4 and IRAK1 distributions had a rise and fall shape, suggesting the assembly of MyD88 was a rate-limiting step requisite for the recruitment of IRAK4 and IRAK1. IRAK4 had a distribution that peaked at ~8 s, and an average recruitment time of 14.4 ± 12.3 s (Fig. 4F, mean ± SD, n = 478 IRAK4 recruitment events). In contrast, IRAK1 had a distribution with a peak at ~18 s and an average recruitment time of 41.3 ± 37.5 s (Fig. 4F, mean ± SD, n = 171 IRAK1 recruitment events). The broad distribution we observed in the IRAK1 distribution of recruitment times possibly reflects that, in addition to the assembly of MyD88, the assembly of IRAK4 serves as a second rate-limiting step. Collectively, we conclude that Myddosomes are an inducible protein complex that forms through the sequential assembly of MyD88, followed by IRAK4 and then IRAK1.

### Myddosomes are highly stable molecular assemblies that do not exchange with uncomplexed MyD88, IRAK4, or IRAK1

Our analysis revealed that the small assemblies of MyD88 were unstable and could disassemble or dissociate from the plasma membrane. In contrast, the larger assemblies of MyD88 showed increased stability and had longer lifetimes at the plasma membrane. The stability of larger complexes could be due to the stable incorporation of components into the Myddosome complex. Alternatively, the Myddosome complex could be kinetically stable, but could still dynamically exchange with uncomplexed MyD88 and IRAK4/1. To distinguish between these possibilities, we applied fluorescent recovery after photobleaching to analyze the assembled Myddosomes.

EL4 cells endogenously expressing MyD88-GFP were incubated with IL1β-functionalized SLBs for 15 mins to allow Myddosomes to form. We selected and photobleached large MyD88-GFP puncta. The photobleached MyD88 puncta did not recover over the experimental time course (60 s post-bleaching, Fig. 5A and Movie S5). We applied the same analysis to EL4 cells endogenously expressing IRAK4-mScarlet and IRAK1-mScarlet. Similar to MyD88-GFP, IRAK4 and IRAK1 puncta did not recover after photobleaching (Fig. 5B-C). Our FRAP analysis of multiple Myddosomes revealed that the core components of MyD88, IRAK4, and IRAK4 do not undergo dynamic exchange (Fig. 5D-F). Thus, these results argue that the Myddosome is a highly stable macromolecular structure with no measurable molecular turnover.

**Figure 5.**
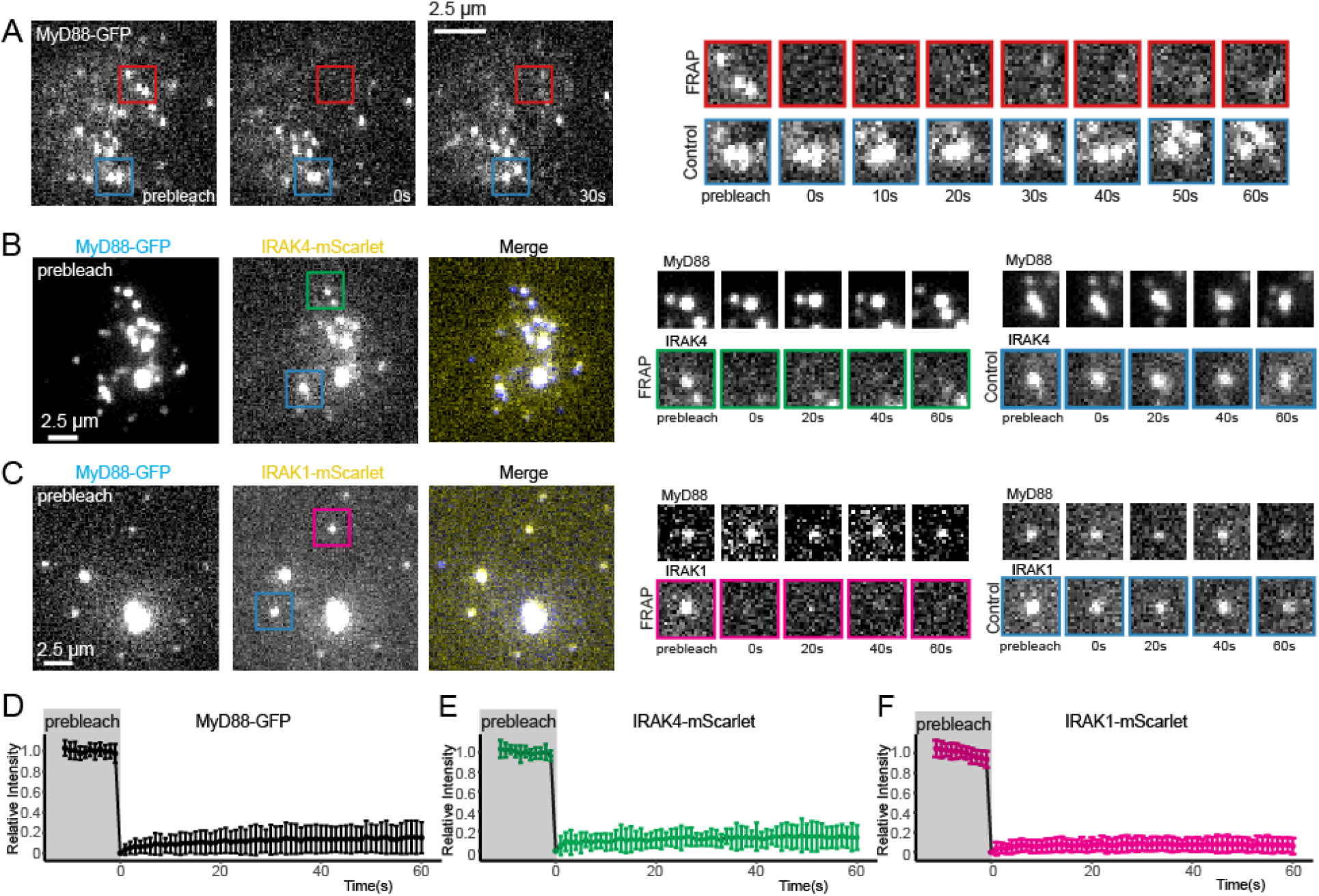
Myddosomes are highly stable molecular assemblies that do not exchange with uncomplexed MyD88, IRAK4, or IRAK1. **A) TIRF images of a MyD88-GFP expressing cell before (prebleach, left), immediately after (0 s, middle) and 30 s after (right) photobleaching**. The red box indicates the position of the Myddosomes on which the FRAP beam was focused. The blue box indicates unbleached control Myddosomes. Right panel, cropped time series of the photobleached and control Myddosomes. **B-C) TIRF images of MyD88-GFP/IRAK4-mScarlet (B) and MyD88-GFP/IRAK4-mScarlet (C) expressing cell before photobleaching.** The red box indicates the position of the IRAK4-mScarlet labelled Myddosome on which the FRAP beam was focused. The blue box indicates unbleached control. Right panels, cropped time series of the photobleached and control IRAK4-mScarlet (B) or IRAK1-mScarlet (C) labelled Myddosome. Scale bar (A-C), 2.5 μm. **(D-F) Normalised recovery curves for MyD88 (n = 61 cells), IRAK4 (n = 22 cells) and IRAK1 (n = 53 cells)**. All yielded a maximal recovery of <20% during the 60 s of observation.

### IRAK4 knockout leads to super MyD88 oligomers

To investigate how MyD88 assembly is regulated by IRAK4/1, we used CRISPR/Cas9 to generate KO cell lines (Fig. S6A). In agreement with previous studies (DeFelice et al., 2019; Suzuki et al., 2002), we found that IRAK4 and IRAK1 KO EL4 cells could not activate NF-kB signaling when stimulated with IL1β (Fig. S6B). We assayed the temporal dynamics of MyD88-GFP puncta in the IRAK4 and IRAK1 KO cell lines. We observed that IL1β stimulation induced the formation of MyD88-GFP puncta in both IRAK4 and IRAK1 KO cell lines (Fig. 6A and Movie S6). This observation suggests that the loss of IRAK4 and IRAK1 does not inhibit the recruitment and oligomerization of MyD88 at activated IL1Rs. However, we did observe that IRAK4 KO cells formed larger (e.g. brighter) MyD88-GFP assemblies. The intensity of MyD88 puncta in IRAK4 KO cells suggests they contain a greater stoichiometry of MyD88 than the 6-8 MyD88s found in purified Myddosome complexes (Lin et al., 2010) (Fig. 6A). Taken together, we argue that IRAK4 functions as a regulator for MyD88 oligomer size, and thus IRAK4 recruitment limits the oligomerization of MyD88.

**Figure 6.**
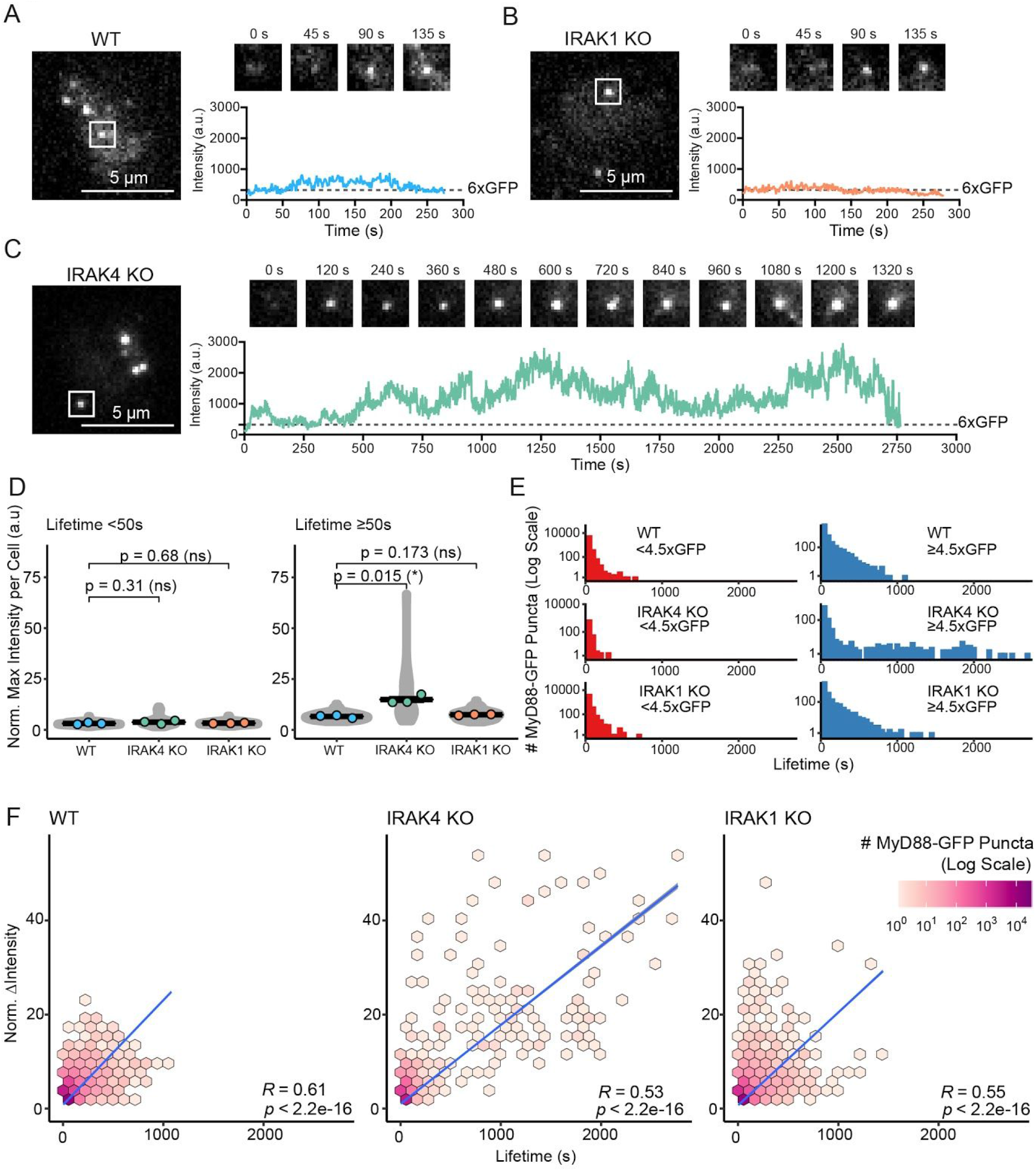
IRAK4 knockout leads to super MyD88 oligomers. We generated IRAK4 and IRAK1 KO cell lines in EL4-MyD88-GFP cells. MyD88-GFP dynamics were assayed using TIRF microscopy. **A-C) TIRF images of MyD88-GFP in EL4 WT (A), IRAK1 KO (B), and IRAK4 KO (C) cells.** Time-series TIRF images from the ROI (white box) showing representative MyD88 puncta. A fluorescence-intensity time trace from each time series is shown below. **D) Long-lived MyD88 puncta have increased maximum fluorescence intensities in IRAK4 KO cells.** Quantification of the maximum intensity of MyD88-GFP puncta per cell with lifetimes of <50 or ≥50 s. Violin plots show the distribution of individual cell measurements. Colored dots superimposed on violin plots correspond to the average value in the independent experiments (n = 3 experimental replicates, encompasses measurements from 10-47 cells). Bars represent mean ± SEM. P-values were calculated using an unpaired Student’s t-test. **E) Larger MyD88-GFP oligomers in IRAK4 KO cells show enhanced lifetime.** Distribution of lifetimes for tracked MyD88-GFP puncta with a maximum intensity of < or ≥4.5x GFP in WT (n = 73,180 for <4.5x GFP and n= 24,631 for ≥4.5x GFP), IRAK4 (n = 18,478 for <4.5x GFP and n= 9,795 for ≥4.5x GFP), and IRAK1 (n = 63,382 for <4.5x GFP and n = 19,735 for ≥4.5x GFP) KO cells. MyD88-GFP puncta lifetimes collated from three experimental replicates. **F) MyD88-GFP puncta in IRAK4 KO cells have increased growth in intensity and longer lifetimes.** 2D histogram of MyD88-GFP puncta lifetime versus change in fluorescent intensity for WT (n = 64,149), IRAK4 KO (n = 13,960), and IRAK4 KO (n = 54,551) cells. Linear fit is shown as a blue line. The coefficient used is Spearman’s rank correlation coefficient.

To examine this possibility, we assessed the distribution of MyD88 oligomer sizes in IRAK4 and IRAK1 KO cell lines. The proportion of larger MyD88 oligomers (i.e. ≥4.5xGFP, Fig. 3B) per cell in IRAK4 and IRAK1 KO cell lines was equivalent to that of WT EL4 cells (Fig. S6D-E). The size of MyD88-GFP puncta with lifetimes of <50 s was equivalent with no statistically significant difference between wild-type and KO cells (MyD88-GFP sizes were 3.12x for WT, 3.78x for IRAK4 KO, and 3.26x for IRAK1 KO, Fig. 6D). In contrast, MyD88 puncta with lifetimes of ≥50 s were twice as large in IRAK4 KO cells compared to both wild-type and IRAK1 KO cells (MyD88-GPFs sizes were 6.58x for WT, 14.9x for IRAK4 KO, and 7.40x for IRAK1 KO, n = 3 experimental replicates for each cell line, Fig 6D). Smaller MyD88-GFP multimers had a similar lifetime distribution across wild-type as well as IRAK4 and IRAK11 KO cell lines However, MyD88-GFP puncta ≥4.5xGFP had extended lifetimes in the IRAK4 KO cells, with several that could be tracked for over 30 mins (Fig. 6E).

In IRAK4 KO cells, we observed stable MyD88-GFP puncta that could be tracked for over 30 mins (Fig. 6C). Over this time these puncta continued to increase in fluorescence intensity (Fig. 6A). This suggested that MyD88 polymerization was unregulated, and led to the formation of super MyD88 oligomers, far greater in size to those that form in wild-type. We then examined the correlation between growth in intensity and lifetime. Similar to our previous analysis (Fig. 3F), we measured a positive correlation between an increase in intensity and the lifetime for MyD88-GFP tracks for wild-type and both the IRAK4 and IRAK1 KO cell lines (R ≥ 0.55 across all wild-type and KO cell lines, Fig. 6F). However, in IRAK4 KO cells, we observed that a portion (n = 110 out of n = 28,273 total MyD88-GFP puncta) of very long-lived events (1000 s) had larger changes in intensity (> 20x GFP) that were not observed in wild type cells (Fig. 6F). This analysis revealed that the greater lifetimes of MyD88-GFP puncta in the IRAK4 KO background correlated with increased growth in intensity (R = 0.55). Therefore, we concluded that the longer lifetime of MyD88-GFP puncta in the IRAK4 KO cells was due to increased MyD88 oligomerization. We suggest that in the absence of IRAK4, MyD88 forms super-assemblies that contain a greater copy number of MyD88 than those observed in previous structural studies (Lin et al., 2010).

We concluded that the absence of IRAK4 leads to unchecked MyD88 assembly. This, in turn, results in larger MyD88 oligomers at the cell surface. This finding suggests that IRAK4 controls MyD88 growth and size (Fig. S7).

## Discussion

The discovery of innate immune receptors and SMOCs argued that macromolecular assembly, in addition to enzymatic activity, can transduce intracellular signaling. However, how does the dynamic process of oligomerization transmit a biochemical signal? To address this question we developed a live-cell assay to investigate the assembly kinetics of Myddosome SMOCs in live cells. We found that Myddosome assembly is dependent on the formation of MyD88 oligomers of a critical size. The formation of large MyD88 oligomers functions as a biochemical threshold that is overcome to activate downstream signaling effectors IRAK4 and IRAK1. Interestingly, MyD88 oligomer size is sensed and controlled by IRAK4 (Fig. S7). Given that multiple innate immune receptors utilize SMOCs, similar mechanistic principles might operate in other innate immune signaling pathways.

### Nucleation and assembly of MyD88 oligomers is reversible and imposes a time delay on signal transduction

The instability of small MyD88 oligomers potentially serves as a safety switch that prevents MyD88 signalling in the absence of stimuli. Relatively few nucleated MyD88 oligomers transitioned to larger signaling competent oligomers (Fig. 3C). The low probability of small oligomers transitioning to a large signalling competent MyD88 oligomers creates a time delay between receptor activation and signal transduction. This could ensure the cells only activate from sustained TLR/IL1R activation resulting from a persistent microbial or environmental threat to the host. This might prevent harmful physiological consequences of auto-activation. Notably, the oncogenic L259P MyD88 mutation has an increased propensity to oligomerize (O’Carroll et al., 2018). This mutation results in sustained NF-kB signaling in the absence of TLR/IL1R stimulation and is a driver mutation in particular B cell lymphomas (Ngo et al., 2011).

We argue that the necessity to suppress auto-activation as well as be reactive to TLR/IL1R activation constrains MyD88 self-assembly. Consistent with this argument, IL1R activation induces the recruitment and self-assembly of MyD88 at the plasma membrane (Fig. 2C). MyD88 oligomerization is initially reversible, and small oligomers (less than 4 MyD88s) are kinetically unstable (Fig. 3D and Fig. S7). Smaller MyD88 oligomers failed to recruit IRAK1 and IRAK4, suggesting limited signaling output (Fig. 4E), and not every interaction between MyD88 and IL1R leads to productive signalling.

### The formation of larger, stable MyD88 oligomers is a decision making step in the TLR/IL1 signaling pathway

Many SMOCs are composed of DD-containing proteins, and structural studies have revealed these effectors can form helical oligomeric complexes (Ferrao and Wu, 2012). Like TLR/IL1Rs and MyD88, many innate immune receptors and their binding effectors do not contain enzymatic activity, but co-assemble and activate downstream enzymes. Our data, coupled with structural studies, support a general mechanism where DD polymer size can create thresholds for triggering the next step of a signalling pathway (Wu, 2013). Compared to smaller oligomers, we found that larger MyD88 oligomers consisting of greater than 4 MyD88s had increased lifetimes (Fig. 3D), and were more likely to recruit IRAK4 and IRAK1 (Fig. 4C-E). We argue that the assembly of larger MyD88 oligomers that can co-assemble with IRAK4 and IRAK1 are a kinetic barrier to signal activation. In this manner, MyD88 oligomer size acts as a physical threshold that must be reached to activate downstream signaling reactions.

Several lines of evidence indicate that a mechanism of oligomer size-dependent thresholding could be operating at inflammasome and TNF receptors. Structural studies have shown that small oligomers of AIM2 or NLRP3 receptors cap larger signalling adaptor ASC oligomers (Lu et al., 2014). This suggests that AIM2/NLRP3 receptors oligomers assembled first and upon reaching a requisite size ASC assembly is triggered. Likewise, the TNF receptor signalling adaptor FADD forms an oligomeric complex at the base of Caspase-8 filaments. Thus, it is likely a required FADD oligomer size is necessary to nucleate Caspase-8 assembly and activation leading to regulated cell death (Fu et al., 2016). Thus, inflammasome receptors and TNF receptor adaptor FADD could function in an equivalent biochemical manner to MyD88.

### Myddosomes assemble on demand with an ordered molecular choreography

Consistent with previous studies (Bonham et al., 2014), we have shown that assembled Myddosomes are only detected after stimulation. However, in comparison to these studies that used immunoprecipitation and Western blot analysis, our novel live-cell image analysis gives the kinetics and precise molecular choreography of Myddosome assembly. We directly measured the sequential assembly of MyD88, IRAK4, and IRAK1 into Myddosomes that occurred over ~1 min time scale (Fig. 5F). Microscopy and biochemical analysis have suggested that oligomers of MyD88 might be present in the cytosol before TLR/IL1R activation (Moncrieffe et al., 2020). While our data does not exclude the presence of preassembled MyD88 oligomers, we found MyD88 oligomerization was inducible and preceded IRAK4/1 recruitment (Fig 3A and Fig. 4A, B), and only larger MyD88 oligomers co-assembled with IRAK4 and IRAK1 (Fig. 4C-D). If preassembled MyD88 oligomers exist, we argue they are not in complex with IRAK1 and IRAK4.

It is important to note that the Myddosome is a signalling complex utilized by nearly all members of the TLR/IL1R superfamily (Gay et al., 2014; O’Neill and Bowie, 2007). Whether different TLRs/IL1R receptors have increased or decreased affinity to MyD88 and therefore can enhance or slow the kinetics of MyD88 oligomerization remains unknown. Furthermore, there are differences in Myddosome composition between TLRs and IL1Rs. Unlike IL1Rs, many TLRs require the TIR domain sorting adapter TIRAP to signal (Horng et al., 2002). TIRAP can co-assemble with TLRs and MyD88, via heterotypic TIR domain interaction, into oligomeric macromolecular complexes (Ve et al., 2017). TIRAP also directs Myddosome formation to membranes enriched in phosphoinositides (Kagan and Medzhitov, 2006). While it is clear that TIRAP has a role in spatially regulating Myddosome signaling, whether or not TIRAP potentiates MyD88 oligomerization to overcome the kinetic bottleneck of forming stable oligomers remains unclear. Future studies are needed to determine whether the kinetics of Myddosome assembly vary across the TLR/IL1R superfamily or in the presence of TIRAP.

### Myddosome stability might be advantageous to signal transduction

The induction of inflammation is a required step for the initiation of a complete immune response, therefore the high stability of SMOCs could be a critical biophysical feature that ensures cells activate a full response. We found by FRAP that once fully formed, Myddosome components do not kinetically exchange (Fig. 5). It is currently unknown whether other DD higher-order assemblies have similar kinetic stability. However, the fact that DD complexes such as the Fas-FADD, FADD-Caspase8, and PIDDosome have highly ordered quaternary structures (Fu et al., 2016; Park et al., 2007; Wang et al., 2010), where subunits have multiple interaction interfaces, suggest a similar intrinsic stability. The prevalence of these complexes in immune signalling pathways suggests this stability might be advantageous to signal transduction. A low dissociation rate might increase the timeframe that downstream reactions, such as the activation of TRAF proteins and ubiquitin ligases (Deng et al., 2000), can be achieved.

### IRAK4 regulates MyD88 oligomer size during signal transduction

Structural studies on DD superfamily signalling protein have revealed complexes with defined stoichiometric ratios of effectors as well as open-ended filamentous structure. Like the Myddosome, the Fas-FADD complexes have defined ratios of 5:5 (Wang et al., 2010). However, how can this be reconciled with DD proteins, such as MyD88, and FADD forming open-ended filaments (Fu et al., 2016; Moncrieffe et al., 2020)? In this study, we observed that loss of IRAK4 results in unchecked MyD88 oligomerization in response to IL1 stimulation (Fig. 6). Given the sequential recruitment of MyD88 and IRAK4 to the cell surface (Fig. 5F), this raises the possibility that IRAK4 biochemically senses a specific size of MyD88 oligomer and restricts further assembly by physically capping the growing end of the MyD88 filament (Fig. S7). *In vitro* studies have observed the dissolution of MyD88 DD filaments into smaller filaments when incubated with IRAK4 DD (Moncrieffe et al., 2020). One possibility is that IRAK4 regulates the size of the MyD88 oligomers via heterotypic DD interactions; however, the precise mechanism remains unknown.

Whole classes of effectors control and regulate cytoskeletal polymer size and growth (Mohapatra et al., 2016). These effectors are critical for the spatial and temporal control of actin and microtubule polymers in diverse cellular processes, such as motility and cell division. We speculate that analogous biochemical effectors, to those discovered in cytoskeletal systems, could regulate the dynamics of SMOC polymers. Regulators of SMOC size and assembly, such as IRAK4, could be critical to building precise signal thresholds for cellular activation.

### Conclusion

We conclude that MyD88 oligomerization is a decision-making step in Myddosome signaling. We propose that the macromolecular assembly of proteins in itself can conceptually be considered a signal transduction step, analogous to phosphorylation in many signalling pathways. Beyond TLR/IL1Rs, multiple innate immune signaling pathways, such as inflammasome receptors, RIG-1 DNA sensors, and TNF receptors, have an equivalent biochemical architecture that consists of receptors and signaling adaptor that self-assemble (Kagan et al., 2014). These diverse receptor systems possibly transduce signals with a similar molecular choreography that begins with the formation of unstable small oligomers that mature into stable larger oligomers that in turn activate downstream signaling. Thus, stepwise assembly, like we have found here for the Myddosome, is likely to be a fundamental feature of SMOC signaling pathways. The study of these diverse innate immune receptors with high spatial-temporal resolution microscopy will lead to a deeper understanding of how protein oligomerization functions in signal transduction.

## Acknowledgments

We thank all members of the Taylor Lab for reagents, experimental advice, and feedback on the manuscript. We thank Luke Larvis (Howard Hughes Medical Institute) for providing JF dyes. We thank the staff at the Advanced Medical Bioimaging Core Facility, Charité Universitätsmedizin Berlin, for help with FRAP data acquisition. We thank Kabir Husain (University of Chicago) for providing image analysis scripts, advice on image analysis and comments on the manuscript. We thank Enfu Hui (University of California San Diego), Lillian Fritz-Laylin (University of Massachusetts Amherst), Olivia Majer (Max Planck Institute for Infection Biology), Elena Levashima (MPIIB) and Helge Ewers (Free University of Berlin) for critical reading of the manuscript. We thank Arturo Zychlinsky (MPIIB) and Scott Dawson (University of California Davis) for comments and editing of the manuscript. This work was supported by the Max Planck Society, and by an International Max Planck Research Student fellowship to Rafael Deliz-Aguirre.

## Methods

### Cell Culture

EL4.NOB1 WT (ECACC, and referred to as EL4 in the paper) and gene-edited cells were grown in RPMI (Thermo Fisher Scientific) with 10% FBS (Biozol) supplemented with 2 mM L-glutamine. EL4 cultures were maintained at a cell density of 0.1-0.5×10^6^ cells/ml in 5% CO_2_, 37°C. HEK-293T cells (ATCC collection) were grown in DMEM (Thermo Fisher Scientific) supplemented with 2 mM L-glutamine and 10% FBS. All cells were determined to be negative for mycoplasma using the MycoAlert detection kit (Lonza).

### Homology-directed repair (HDR) DNA template design for CRISPR/Cas9 endogenous labeling

Plasmid DNA repair templates were designed using a pMK (Life Technologies, Carlsbad, CA) vector backbone. Silent mutations were included in the homology arms to remove sgRNA target sites and avoid Cas9 cleavage of the repair template. Homology arms were amplified from EL4 Genomic DNA, and assembled with DNA fragments encoding a fluorescent protein tag (e.g., mEGFP or mScarlet-i (Bindels et al., 2017)) and pMK plasmid backbone using Gibson Assembly. All HDR template plasmids were sequence verified. Full details of the HDR DNA template plasmid construction are given below (full sequences of the HDR templates given in Table S7).

### pMK-MyD88-mEGFP-HDR

5’ and 3’ homology arms were designed from the mouse *MyD88* gene (ENSMUSG00000032508) covering a distance of 1015 bps and 1069 bps either side of the TGA stop codon. mEGFP was inserted between these homology arms and fused to the MyD88 C-terminus via a 3x(Gly-Gly-Ser) linker.

### pMK-IRAK4-mScarlet-i-HDR

5’ and 3’ homology arms were designed from the mouse *Irak1* gene (ENSMUSG00000059883) covering a distance of 702 bps and 722 bps either side of the TAA stop codon. mScarlet-i was inserted between these homology arms and fused to the IRAK4 C-terminus via a 3x(Gly-Gly-Ser) linker.

### pMK-IRAK1-mScarlet-i-2A-PuroR-HDR

5’ and 3’ homology arms were designed from the mouse *Irak1* gene (ENSMUSG00000031392) covering a distance of 2251 bps and 764 bps either side of the TGA stop codon. A mScarlet-i-2A-PuroR cassette was inserted between these homology arms and fused to the IRAK1 C-terminus via a 3x(Gly-Gly-Ser) linker.

### Generation of CRISPR/Cas9 sgRNA vectors for endogenous labelling of MyD88, IRAK4 and IRAK1

Single-guide RNAs (sgRNA) targeting +/− 50 bps of the C-terminus stop codon of MyD88, IRAK4 and IRAK1 were designed using the web-based Benchling CRISPR design tool. sgRNAs were selected for each target (Table S1), and complementary oligonucleotides designed to be ligated into Bbs1 digested *Streptococcus pyogenes* Cas9 and chimeric guide RNA expression plasmid pX330, (pX330-U6-Chimeric_BB-CBh-hSpCas9, Addgene #42230). sgRNA oligonucleotides were ordered from Integrated DNA Technologies (IDT). Complementary sgRNA oligonucleotides were 5’ phosphorylated with T4 Polynucleotide kinase, annealed, ligated into Bbs1 digested pX330 using Quick Ligase (NEB). pX330 plasmids were transformed into *NEB Stable* competent cells. All sgRNA pX330 plasmids were sequence verified.

### Generation of CRISPR/Cas9 Engineered Cell Lines

EL4 cells were electroporated with pX330 Cas9/gRNA expressing vector and the pMK vector encoding the HDR template with the Neon Transfection System. EL4 cells were electroporated with the following conditions: voltage (1080 V), width (50 ms), number of pulses (one). For single editing of the MyD88 gene locus, 1.5μg total of MyD88 sgRNA-Cas9 and MyD88-GFP HDR template plasmids (in equal molar ratio) were electroporated into 2×10^7^ cells per ml for a 10μl reaction with buffer R according to the manufacturer’s protocol. Cells were plated to 24-well plates in RPMI culture medium without antibiotics and expanded for seven days.

Monoclonal cell lines were generated by fluorescence-activated cell sorting (FACS). Cells were sorted using BD FACS Aria II at Deutsches Rheuma-Forschungszentrum Berlin, Flow Cytometry Core Facility. To isolate gene-edited EL4 cells, we performed a bulk sorting of GFP positive cells (Fig. S1B). This population was expanded and single cell lines sorted onto 96-well plates containing culture medium with 15% EL4.NOB-1 conditioned RPMI medium. The same strategy was applied for double editing of MyD88/IRAK4 or MyD88/IRAK1 gene loci. 1.5 μg of sgRNA-Cas9 and HDR template plasmids (in equal molar ratio) were electroporated simultaneously. For the selection of IRAK1 edited events, 1.5 μg/ml Puromycin was added to the cell culture medium 24h after electroporation. EL4 cells were selected in Puromycin for 48hr.

Monoclonal cell lines were verified using PCR, sequencing, western blot analysis, and microscopy (Fig. S1A-D and S4B,C). First, genomic DNA was isolated from selected monoclonal cell lines using QuickExtract DNA Extraction Solution (Epicentre). To test for gene editing and positional insertion of mGFP/mScarlet-I cassette, PCR primers were designed to amplify a DNA fragment that contained the junctions between mGFP/mScarlet-I open reading frame, the 3’ or 5’ homology arm and the gene locus (see Table S2-4 for primer sequences). To check single-cell clones for homozygosity, we designed PCR primers that amplified a fragment containing mGFP/mScarlet-I cassette, the entire 3’ or 5’ homology arms and the junction between the homology arms and the gene locus (see Table S2-4 and Fig. S1C). PCR products were analysed on a 0.8-1% agarose gel. Homozygosity was detected by the presence of a single high molecular weight DNA band (Fig. S1C). Gel fragments of homozygous clones were extracted using Monarch Nucleic Acid Purification Kits (NEB) and submitted for Sanger Sequencing. To confirm the presence of mEGFP/mScarlet-I fusion protein of the correct molecular weight, CRISPR/Cas9 edited cell clones were analysed by western blot. Lysates were blotted with antibodies specific for the target protein, and then stripped and re-probed with antibodies specific for GFP or mScarlet-I (Fig. S1D, S4B,C, full-length Western blot shown in Fig. S7, S8 and S9). Finally, all cell clones were checked by microscopy for correct localisation of fluorescent signals.

### Generation of CRISPR/Cas9 sgRNA IRAK4 and IRAK1 knockout vectors

Single-guide RNAs (sgRNA) targeting the first coding exon of the N-terminus of IRAK4 and IRAK1 were designed using the web-based Benchling CRISPR design tool. sgRNAs were selected for each target, and complementary oligonucleotides designed to be ligated into Bbs1 digested *Streptococcus pyogenes* Cas9 and chimeric guide RNA expression plasmid pX459v2.0, (pX459v2.0-pSpCas9(BB)-2A-Puro, Addgene #62988) or pX459v2.0-HypaCas9 (pX459v2.0-HypaCas9-2A-Puro, Addgene #108294). sgRNA oligonucleotides were ordered from Integrated DNA Technologies (IDT).

Complementary sgRNA oligonucleotides were 5’ phosphorylated with T4 Polynucleotide kinase, annealed, ligated into Bbs1 digested pX459v2.0 using Quick Ligase (NEB). pX330 plasmids were transformed into *NEB Stable* competent cells. All sgRNA pX459v2.0 plasmids were sequence verified.

### Homology-directed repair (HDR) DNA template design for CRISPR/Cas9 generation of IRAK1 and IRAK4 KO cells

Plasmids DNA repair templates to generate KO cell lines were designed in the same way as described above. The homology arms were assembled with DNA fragments encoding a blasticidin resistant cassette followed by 3xSTOP codons and +1nt (to induce a downstream frameshift) into pMK plasmid backbone using Gibson Assembly. All HDR KO template plasmids were sequence verified. Full details of the HDR DNA template plasmid construction are given below. Full sequences of HDR templates are reported in Table S7.

### pMK-IRAK4-KO-BlastR-3xStop-HDRtemp

5’ and 3’ homology arms were designed from the mouse *Irak4* gene (ENSMUSG00000059883) covering a distance of 254 bps and 657 bps either side of the ATG start codon. Blasticidin resistance cassette with 3’ 3xSTOP codons plus 1 nt was inserted between these homology arms.

### pMK-IRAK1-KO-BlastR-3xStop-HDRtemp

5’ and 3’ homology arms were designed from the mouse *Irak1* gene (ENSMUSG00000031392) covering a distance of 896 bps and 874 bps either side of the ATG start codon. Blasticidin resistance cassette with 3’ 3xSTOP codons plus 1 nt between these homology arms.

### Generation of CRISPR/Cas9 IRAK1 and IRAK4 knockout Cell Lines

Two methods were used to generate IRAK1/IRAK4 KO cell lines. In the first methods, EL4-MyD88-GFP cells were electroporated with pX459v2.0 Cas9/gRNA. 24 hrs after electroporation the cells were selected in puromycin for 3 days. After selection, cells were single cells sorted into 96 well plates. Isolated clones were then screened (see below for details). We found this method inefficient with many clones still WT. Only one IRAK1 KO clone was isolated using this method (Fig. S6B, clone 1).

To enrich from KO cells we developed a second method that used HDR templates to insert a blasticidin cassette followed by 3x STOP codons. In this method EL4-MyD88-GFP cells were electroporated with the pX459v2.0 Cas9/gRNA and a pMK vector encoding the KO-HDR template with the Neon Transfection System (see above for conditions). Electroporation cells were maintained in complete RPMI media for three days after electroporation. On the fourth day cells were split into RPMI media containing blasticidin (6 μg/ml). Cells were selected for 7-14 days with blasticidin and then sorted in 96 well plates to select single cell clones.

Monoclonal KO cell lines were verified using PCR, sequencing, and western blot analysis, (Fig. S1A-D and S4B,C). First, genomic DNA was isolated from selected monoclonal cell lines using QuickExtract DNA Extraction Solution (Epicentre). Primers specific to the blasticidin resistance cassette and IRAK1/IRAK4 gene loci were used to verify the insertion (see Table S5 and S6 for IRAK4 and IRAK1 respectively). We designed a second set of PCR primers that amplified a fragment containing the blasticidin cassette, the entire 3’ or 5’ homology arms, and the junction between the homology arms and the gene locus (see Table S5 and S6). PCR products were analysed on a 0.8-1% agarose gel, and gel fragments of clones were extracted using Monarch Nucleic Acid Purification Kits (NEB) and submitted for Sanger Sequencing. We found only homozygous Blasticidin insertion clones for the KO of IRAK1. In contrast with IRAK4 we found only heterozygous clones; however sequencing confirmed the presence of an insertion in the second allele resulting in a frameshift. All clones analysis by Western blot analysis to be KO for IRAK4 or IRAK1 (Fig. S6A,B and Fig S8-9 for full-length Western blots).

### Imaging Chambers and Supported Lipid Bilayers

SLBs were prepared using a previously published method (Taylor et al., 2017). Briefly, Phospholipid mixtures consisting of 97.5% mol 1-palmitoyl-2-oleoyl-*sn*-glycero-3-phosphocholine (POPC), 2% mol 1,2-dioleoyl-sn-glycero-3-[(N-(5-amino-1-carboxypentyl)iminodiacetic acid)succinyl] (ammonium salt) (DGS-NTA) and 0.5% mol 1,2-dioleoyl-sn-glycero-3-phosphoethanolamine-N-[methoxy(polyethylene glycol)-5000] (PE-PEG5000) were mixed in glass round-bottom flasks and dried down with a rotary evaporator. All lipids used were purchased from Avanti Polar Lipids. Dried lipids were placed under vacuum for 2 hrs to remove trace chloroform and resuspended in PBS. Small unilamellar vesicles were produced by several freeze-thaw cycles. Once the suspension had cleared, the lipids were spun in a benchtop ultracentrifuge at 65,000xg for 45 min and kept at 4°C for up to five days.

Supported lipid bilayers were formed in 96-well glass bottom plates (Matrical) or set up on coverslips (25 mm diameter, No. 1.5 H, Marienfeld-Superior) mounted in Attofluor chamber (Thermo Fisher). 96-well plates were cleaned for 30 min with a 5% Hellmanex solution containing 10% isopropanol heated to 50°C, then incubated with 5% Hellmanex solution for 1 hour at 50°C, followed by extensive washing with pure water. 96-well plates were dried with nitrogen and sealed until needed. To prepare SLB on 96-well plates, individual wells were cut out and base etched for 15 min with 5 M KOH and then washed with water and finally PBS. Coverslips were washed in acetone and ethanol in an ultrasonic cleaner, before rinsing extensively in water. Coverslips were then cleaned with a solution of KOH and hydrogen peroxide for 10 mins, followed by extensive washing in water. Finally coverslips were cleaned with a solution of 6% HCl (V/V) and 6.3% (V/V) hydrogen peroxide. Cleaned coverslips were stored in water before being used for SLBs formation and microscopy.

To form SLBs, SUVs suspension were deposited in each well or coverslip and allowed to form for 1 hr. We found that SUVs suspension containing 0.5% mol PE-PEG5000 formed best at 45°C. After 1 hr, wells were washed extensively with PBS. SLBs were incubated for 15 min with HEPES buffered saline (HBS: 20 mM HEPES, 135 mM NaCl, 4 mM KCl, 10 mM glucose, 1 mM CaCl_2_, 0.5 mM MgCl_2_) with 5 mM NiCl_2_ to charge the DGS-NTA lipid with nickel. The SLBs were then washed in HBS containing 1% BSA to block the surface and minimize non-specific protein adsorption. After blocking, the SLBs were functionalized by incubation for 1 hr with his-tagged proteins. The labeling solution was then washed out and each well was completely filled with HBS with 1% BSA. For SLBs set up on 96-well plates the total well volume was 625 μl (manufacturers specifications), and 525 μl was removed leaving 100 μl of HBS 1% BSA in each well.

### Protein Expression, Purification and Labeling

To functionalize membranes with active mouse IL1β, we created a protein linker that could tether the mature IL1β cytokine to SLBs. To aid in solubility and expression, we designed this tether not to be directly fused to mature IL1β on the same peptide chain. We used a SpycatcherV2 domain to covalently link this tether to recombinant mature mouse IL1β expressed with a c-terminus SpytagV2 peptide (AHIVMVDAYKPTK). Spycatcher is an engineered protein domain derived from the *S. pyogenes* CnaB2 domain that is able to form an isopeptide bond to the SpyTag peptide (Keeble et al., 2017). To construct this protein tether, we created a codon-optimised HaloTag and SpycatcherV2 open reading frames and ordered them as gBlocks from IDT. To enhance solubility and expression of this fusion protein, the HaloTag and Spycatcher open reading frames were separated by a Tencon domain (a high-stability FNIII domain designed through multiple-sequence alignment (Jacobs et al., 2012)). Using Gibson Assembly, gene fragments were cloned into a pET28a vector containing a N-terminal 10x His tag (pET28a-His10-Halo-Tencon-SpycatcherV2). We also created an identical version where mScarlet was substituted for HaloTag (pET28a-His10-mScarlet-i-Tencon-SpycatcherV2). The mature active form of mouse IL1β (aa. 118-169) was codon optimised for E coli expression and ordered as a gBlock from IDT. The gBlock was designed to contain a c-terminal SpyTagV2 connected via a 13aa Glycine Serine linker. We used Gibson Assembly to clone this fragment into pET28a (pET28a-MmIL1β-Spytag).

All proteins were expressed in BL21-DE3 Rosetta *E. coli* (Novagen). The bacterial cell pellets were resuspended in the lysis buffer (20 mM HEPES, 150 mM NaCl with protease inhibitors) and lysed using a French press. To covalently couple His10-Halo/mScarlet-i-Tencon-Spycatcher to MmIL1β-Spytag, the cleared lysates were mixed and incubated with mild agitation for 1 hour at 4 C. To ensure complete spycatcher-spytag conjugation, the lysates were mixed with 2:1 ratio (vol:vol, based on starting bacterial culture volume) of MmIL1β-Spytag to His10-Halo/mScarlet-I-Tencon-Spycatcher. After the conjugation, the His10-Halo/mScarlet-I-Tencon-Spycatcher-IL1β-Spytag was purified by Ni-NTA resin. Conjugation was monitored by mobility shift using SDS-PAGE. After elution, the protein was dialysed into 20 mM HEPES overnight, followed by anion exchange chromatography on a MonoQ column. This was followed by gel filtration over Superdex 200 26/600 into storage buffer (20 mM HEPES, 150 mM NaCl). The protein was snap frozen with the addition of 20% glycerol in liquid nitrogen and placed at −80°C for long-term storage. In text these proteins are referred to as His10-mScarlet-IL1β or His10-Halo-IL1β.

Following purification, the His10-Halo-Tencon-Spycatcher-IL1β-spytag protein was either snap-frozen and stored at −80°C or directly used for HaloTag-labeling. To label the HaloTag, a 2.5x molar excess of JF646-HaloLigand was mixed with the protein and incubated at room temperature for 1 hour followed by an overnight incubation at 4°C. Post labelling, the protein was gel filtered over a Superdex 200 26/600 into storage buffer and snap-frozen with the addition of 20% glycerol in liquid nitrogen and placed in −80°C for storage. The degree of labelling was calculated with a spectrophotometer by comparing 280 nm and 640 nm absorbance (usually 85-95% labeling efficiency was achieved).

### Immunofluorescence analysis of RelA and phospho-p38 localisation

To analyse the nuclear localisation of RelA or phospho-p38 levels in IL1β stimulated EL4 cells, SLB were labelled with fluorescent IL1β, and unlabelled SLBs served as unstimulated controls. On the day of an experiment EL4.NOB1 cells endogenously expressing MyD88-GFP were transferred to serum free media and incubated for 3-4 hrs. Before stimulation, EL4 cells were resuspended in HBS at a final concentration of 3×10^6^ cells per ml. 100 μL of cells (corresponding to 3×10^5^ cells per coverslip chamber) were then applied to each supported membrane. Cells were then incubated at 37 °C for 30-45 min before the addition of an equal well volume of 2x fixative (7% (v/w) PFA with 1% (v/w) Triton X). Cells were fixed for 20 min at room temperature. Cells were then washed with PBS containing 60 mM glycine to quench PFA. Cells were then blocked in PBS 10% (w/v) BSA for 1 hr at room temperature or overnight at 4°C before addition of primary antibody. Fixed cells were labelled overnight with primary antibodies diluted in PBS 10% (w/v) BSA (for anti-RelA: rabbit monoclonal, Cell Signaling Technology #8242, 1:400; for phospho-p38: rabbit monoclonal Cell Signaling Technology #4511, 1:160000). The next day cells were washed 5x in PBS 10% (w/v), and labeled with secondary antibodies (goat anti-rabbit conjugated to Alexa Fluor 555/647,Invitrogen, 1:1000) and FluoTag®-X4 anti-GFP conjugated to Atto488 (Nano Tag Biotechnology, 1:1000 to boost the MyD88-GFP signal) for 1 hr at room temperature or overnight at 4°C. Cells were washed 5x in PBS. In the penultimate PBS wash, cells were labeled for 10 min with DAPI at a labeling concentration of 300 nM.

For confocal imaging, coverslips were mounted on slides using Vectashield (Vectorlabs). Mounted coverslips of stimulated and unstimulated EL4 cells were imaged on a Zeiss Airyscan LSM 880 using a Plan Apo 63x 1.4 NA oil-immersion objective. Fields of view containing multiple cells were selected based on the DAPI and MyD88-GFP channel. Z-stacks of the entire cellular volume were acquired in the DAPI, GFP and Alexa647 channel. The nuclear staining intensity of RelA and phospho-p38 was analysed in Fiji. Z-stacks of the DAPI, GFP, Alexa647 channel were imported into FIJI. Using the DAPI channel 3 Z-planes were selected and a maximum projection of the Z-planes was used to create a new 32-bit image. Maximum projection images were created of the identical Z-planes for the Alex647 channel. The DAPI channel was segmented to identify cell nuclei. The detected nuclear boundaries were used to extract nuclear staining intensity from the Alex647 channel (e.g the RelA or phospho-p38 staining intensity, Fig. 1E and Fig. S1G).

### TIRF-Microscopy data acquisition

Imaging of MyD88-GFP, IRAK recruitment was performed on an inverted microscope (Nikon TiE, Tokyo, Japan) equipped with NIKON fiber launch TIRF illuminator. Illumination was controlled with an laser combiner using the 488, 561 and 640 nm laser lines at approximately 0.35, 0.25 and 0.17 mW laser power respectively (laser power measured after the objective). Fluorescence emission was collected through filters for GFP (525 ± 25 nm), RFP (595 ± 25 nm) and JF646 (700 ± 75 nm). All images were collected using a Nikon Plan Apo 100x 1.4 NA oil-immersion objective that projected onto a Photometrics 95B Prime sCMOS camera with 2×2 binning (calculated pixel size of 150 nm) and a 1.5x magnifying lens. Image acquisition was performed using NIS-elements software. All experiments were performed at 37°C. The microscope stage temperature was maintained using an OKO Labs heated microscope enclosure. Images were acquired between intervals of 1-5 s using exposure times of 70–100 ms.

### Imaging EL4 cells endogenously expressing MyD88-GFP, IRAK4-mScarlet or IRAK1-mScarlet on IL1β functionalized SLBs with TIRF-Microscopy

His10-Halo-JF646-IL1β functionalized SLBs were set up as described above. SLBs were functionalized with 1-5 nM of His10-Halo-JF646-IL1β. To quantify the density of IL1β on the SLB, wells were prepared that were functionalized with identical labelling protein concentration and time, but with different molar ratios of labelled to unlabelled His10-Halo-IL1β. Before application of cells, SLBs were analysed by TIRF microscopy to check formation, mobility and uniformity. Short time series were collected at wells containing a ratio of labelled to unlabelled His10-Halo-IL1β, (e.g. <1 His10-Halo-JF646-IL1β molecule per μm^2^) to calculate ligand densities on the SLB based upon direct single molecule counting. All experiments were performed at IL1 SLB densities of 10 to 30 molecules per μm^2^.

Before each imaging experiment we acquired calibration images using recombinant mEGFP. To image single GFP fluorophore, recombinant purified monomeric EGFP was diluted in HBS and adsorbed to KOH cleaned glass. Single molecules of GFP were imaged using identical microscope acquisition settings to those used for cellular imaging. To image live cells, EL4 cells were pipetted onto supported lipids bilayers functionalized with His10-Halo-JF646-IL1β. EL4 cells expressing only MyD88-GFP were sequentially illuminated for 70-100 ms with 488 nm at a frame interval of 1 s (Fig. 3). EL4 cells expressing MyD88-GFP, IRAK4-mScarlet or IRAK1-mScarlet were sequentially illuminated for 70-100 ms with 488 nm and 100ms with 561nm laser line at a frame interval of 1 s (Fig. 4). Diffraction-limited punctate structures of MyD88-GFP, IRAK4-mScarlet or IRAK1-mScarlet were detected and tracked using the FIJI TrackMate plugin (Tinevez et al., 2017).

### FRAP experiments and data analysis

FRAP experiments were performed at the Advanced Medical BioImaging Core Facility at the Charité, on a NIKON TIRF microscope (Nikon Eclipse Ti-E) with the FRAPPA module. Fluorescent images were acquired with a Nikon Plan Apo 100x 1.4 NA oil-immersion objective and projected on a Photometric Prime 95B sCMOS camera. For FRAP analysis of Myddosomes in gene edited EL4 cells, we prepared SLBs as described above. We labelled SLBs with either IL1β-mScarlet-His10 or IL1β-Halo-JF646-His10 depending whether the photo-bleach experiments were performed in EL4-MyD88-GFP or EL4-MyD88-GFP/IRAK4-mScarlet and EL4-MyD88-GFP/IRAK1-mScarlet gene-edited cell lines. Before the addition of cells to the imaging chamber, we analyzed SLB formation and mobility by visual inspection of the fluorescently labelled IL1β. We prepared EL4 cells for imaging by washing in PBS and resuspending in HBS at a final concentration of 1 x 10^5^ cells per ml. To ensure FRAP analysis of fully assembled Myddosomes, we incubated 1 x 10^4^ EL4 cells with IL1β functionalized SLBs for at least 15 mins before image acquisition. After incubation, we selected EL4 cells containing multiple MyD88-GFP labelled Myddosome for FRAP analysis. The photobleach spot was centered on large stationary Myddosomes or in some cases a cluster of Myddosomes. Images were recorded 10 sec before and 60 sec after photobleaching at a time interval of 1 image per second. The FRAP laser beam was set up accordingly: 6% 488 nm laser power for MyD88-GFP, and 10% 561 nm laser power for IRAK1/IRAK4-mScarlet. All fluorescent images were acquired using an exposure time of 100 ms.

To quantify FRAP recovery we adapted the approach described by Kang et al. (Kang et al., 2015). We determined the integrated intensity of the photobleached region as a function of time. The background intensity was measured from a neighboring region to the photobleached spot and was subtracted from all timepoints. The data was normalized to the pre-bleach intensity using the following equation: 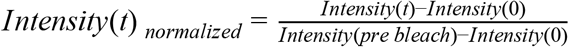, where *Intensity(pre bleach*) is the averaging intensity preceding photobleaching, and *Intensity(0)* is the intensity immediately preceding photobleaching. Measurements from multiple photobleached Myddosomes were averaged, and the standard deviation was calculated (Fig. 5D-F). We conducted three independent experimental replicates on different days for each FRAP experiment (e.g. MyD88-GFP, IRAK1-mScarlet and IRAK4-mScarlet).

## Quantification and Statistical Analysis

All data are expressed as the mean ± the standard deviation (SD) or mean ± the standard error of the mean (SEM), as stated in the figure legends and results. The exact value of n and what n represents (e.g., number of cells, MyD88-GFP puncta, or experimental replicates) is stated in figure legends and results. Means were compared using an unpaired Student’s t-test. Data and scripts used in this study available at 10.5281/zenodo.4012312 and gitlab.com/Marcus_Taylor/myddosome-dynamics-pipeline.

### Quantification and analysis of MyD88-GFP intensity, lifetime and dynamics

To quantify the dynamics of MyD88-GFP assemblies in EL4 cells, we created an image analysis pipeline that ran in Fiji, MATLAB and R. The MyD88-GFP fluorescence channel images were processed in Fiji to remove background intensity using custom-written macros. First, we subtracted a dark frame image (e.g. an image only containing intensity values from current and noise generated by the camera electronics) from each image file. The dark frame image was created by averaging 5000 camera images captured without light exposure and with the same shutter speed as the images. The MyD88-GFP channel images were processed with a median filter (radius 25 pixels) to create an image of the cytosolic background intensity. The resulting median-filtered image of the background intensity was subtracted from the dark frame-subtracted image stack to create a new MyD88-GFP image stack. We quantified the MyD88-GFP signal intensity using this background-subtracted image stack.

Individual cells were identified in the MyD88-GFP fluorescence channel using either a marker-controlled watershed segmentation (implemented with the MorphoLibJ ImageJ plugin (Legland et al., 2016) using a maximum projection of the MyD88-GFP fluorescence channel) or manually. We used the Fiji TrackMate plugin to track the MyD88-GFP particles within each segmented cell. Tracking coordinates generated by TrackMate were imported into MATLAB, and the fluorescence intensity of MyD88-GFP puncta were measured from a 3×3 pixel region. To compute the distribution of single fluorophore intensities, images of single mGFP fluorophores absorbed to glass were processed and analysed identically to MyD88-GFP images. After background subtraction and particle tracking, subsequent analysis was performed in R. We restricted the analysis to MyD88-GFP puncta tracked for three or more frames, to focus our analysis to bona fide MyD88 assemblies nucleating at the plasma membrane.

To analyse the size distribution and stoichiometry of MyD88 multimers, we identified the fluorescence intensity maxima for each tracked MyD88-GFP puncta (Fig. 3B and Fig. S3A). To quantify the size (e.g., the copy number of MyD88-GFPs in a tracked puncta), we divided the fluorescent intensities by the mean intensity of single mEGFP fluorophores (measured before each experiment using the same acquisition setting as cellular data, see above). Structural studies that have identified 6-8 MyD88 monomers in purified Myddosome complexes (Lin et al., 2010). We reasoned that brighter MyD88-GFP diffraction-limited puncta corresponded to larger multimers of MyD88 and therefore fully assembled Myddosomes. To identify large multimers of MyD88, we estimated the fluorescent intensity distribution for diffraction-limited particles containing 6x GFPs. We estimated the fluorescent distribution of 6x GFP to be Gaussian with a mean and variance equal to 6x single GFP fluorophores. Therefore, we used a threshold of ≥4.5x GFP to exclude tracks that consist of MyD88 monomers, dimers or trimers. We calculated this threshold would select >98% of MyD88-GFP tracks containing 6x MyD88-GFP (although smaller assemblies of 4x and 5x would also be selected). We used this intensity threshold to categorise MyD88-GFP puncta as small (e.g., less <4.5x GFP) or large (≥4.5x GFP) MyD88 oligomers. We performed data visualisation (e.g., of density plots of intensity maxima distribution and percentage of categorised MyD88 puncta per cells) using ggplot, a data visualisation package for R.

To examine the growth of tracked MyD88 puncta, we calculated the maximal growth by subtracting the maximal intensity from the starting intensity (Fig. 3F). The change in intensity was normalized by dividing it by the intensity of mEGFP. We performed this analysis on MyD88-GFP tracks that had an initial fluorescent intensity <2.5x the mean intensity of GFP. Manual inspection of the date revealed that this cut-off restricted the analysis to nucleating assemblies of MyD88-GFP, and removed those that bud or split from pre-existing assemblies where the time point of nucleation could not be accurately determined (Fig. S2B). We tested correlation between ΔIntensity and lifetime using Spearman’s rank correlation analysis. The correlation coefficient (R) is reported in the figure legends and text. A 2D histogram of ΔIntensity norm versus lifetime was plotted using ggplot (Fig. 3F and Fig. S3E).

### Analysis of IRAK4 and IRAK1 colocalization with MyD88

In data acquired with EL4 cells expressing MyD88-GFP and IRAK4/IRAK1-mScarlet, the IRAK4-mScarlet and IRAK1-mScarlet fluorescent channel was processed using an identical image processing pipeline that was applied to the MyD88-GFP channel (e.g. background subtraction, see above). The IRAK4 and IRAK1 mScarlet was broken up into minstacks of individual cells. We used the Fiji TrackMate plugin to identify and track IRAK4/1 fluorescent puncta. To quantify colocalization between the MyD88-GFP and IRAK4/1-mScarlet fluorescent channels, we imported the tracking coordinates generated by TrackMate into MATLAB. Using these coordinates, we identified MyD88 and IRAK4/1 particles that colocalized based on two or greater consecutive frames where the tracked coordinates were equal or less than 0.25 μm apart. Using this criteria MyD88 tracked puncta were classified as either positive or negative for IRAK4/1 colocalization.

## Supplementary Movies

**Movie S1**: IL1β tethered to a supported lipid membrane forms clusters that recruit MyD88-GFP (related to Fig. 2). This movie shows an EL4 cell expressing MyD88-GFP (middle panel, green channel in merge) interacting with a supported lipid bilayer functionalized IL1β-JF646 (right panel, magenta channel in merge). The movie illustrates that the IL1β clustering at the cell-supported membrane interface precedes the recruitment and formation of MyD88-GFP puncta at the cell surface. Same cell as shown in Fig. 2B. Scale bar, 5 μm.

**Movie S2:** Single cell analysis of MyD88-GFP puncta dynamics (related to Fig. 3). The movie shows an EL4 cell expressing MyD88-GFP imaged using TIRF microscopy, and illustrates MyD88 puncta tracking and analysis of MyD88-GFP puncta for a single cell. Analysis is updated as the movie progresses. Panels in the movie correspond to the following and follow the same order as Fig. 3. **A)** Movie of EL4 MyD88-GFP imaged under TIRF microscopy with a time interval of 1 second. Overlay of particle track trajectories. Particle trajectory colored coded according to whether the max intensity is ≥4.5x GFP (blue) or <4.5x GFP (red). **B)** Density plot of the maximum fluorescent intensity of tracked MyD88-GFP puncta from the cell in (A) (dark blue curve). Intensity distribution of GFP and 6x GFP multimer (green and light blue, respectively) is shown for comparison. Blue background shade indicates ≥4.5xGFP. **C)** Proportion (%) of MyD88-GFP puncta with a maximum intensity <4.5x GFP (red) or ≥4.5x (blue). **D)** Lifetime distribution of MyD88-GFP puncta. MyD88-GFP lifetime histogram for MyD88-GFP puncta with a maximum intensity of <4.5x GFP (red) or ≥4.5x GFP (blue). The puncta count is shown in log scale. **E)** Proportion (%) of MyD88-GFP puncta with a maximum intensity of ≥4.5x GFP with lifetimes of <50 or ≥50 s. **F**) Correlation between lifetime and intensity growth of MyD88-GFP puncta. 2D histogram of MyD88-GFP puncta lifetime by change in fluorescent intensity. The linear regression line is shown in blue with 95% confidence intervals in gray. The MyD88-GFP puncta count is shown in log scale.

**Movie S3**. IRAK4 recruitment to clusters of MyD88 (related to Fig. 4). This movie shows an EL4 cell expressing MyD88-GFP (left panel, green channel in merge) and IRAK4-mScarlet interacting with an IL1β-functionalized supported lipid membrane. The movie illustrates the nucleation of MyD88-GFP puncta and the recruitment of IRAK4-mScarlet. Same cell as shown in Fig. 4A. Scale bar, 5 μm.

**Movie S4**. IRAK1 recruitment to clusters of MyD88 (related to Fig. 4). This movie shows an EL4 cell expressing MyD88-GFP (left panel, green channel in merge) and IRAK4-mScarlet interacting with an IL1β-functionalized supported lipid membrane. The movie illustrates the nucleation of MyD88-GFP puncta and the recruitment of IRAK4-mScarlet. Same cell as shown in Fig. 4A. Scale bar, 5 μm.

**Movie S5**. FRAP analysis of MyD88-GFP (related to Fig. 5). This movie shows an EL4 cell expressing MyD88-GFP in which a region of interest (red box) is photobleached. The movie illustrates that Myddosomes show no fluorescence intensity recovery after photobleaching. Same cell as shown in Fig. 5A. Scale bar, 5 μm.

**Movie S6.** MyD88-GFP dynamics in WT, IRAK4 KO and IRAK1 KO EL4 cells (related to Fig. 6). This movie shows MyD88-GFP in WT (left panel), IRAK1 KO (middle panel) and IRAK4 KO EL4 cells lines landing on an IL1β-functionalized supported lipid membrane. White box in each panel indicates example cells shown in Fig. 6A. Scale bar, 5 μm.

## Supplementary Tables

**Table S1.** Key resources table

**Table S2.** PCR primers for MyD88 sequence validation

**Table S3.** PCR primers for IRAK4 sequence validation

**Table S4.** PCR primers for IRAK1 sequence validation

**Table S5.** PCR primers for IRAK4 KO sequence validation

**Table S6.** PCR primers for IRAK1 KO sequence validation

**Table S7.** Sequences of the *HDR templates*

---

**Supplementary Figure 1.**
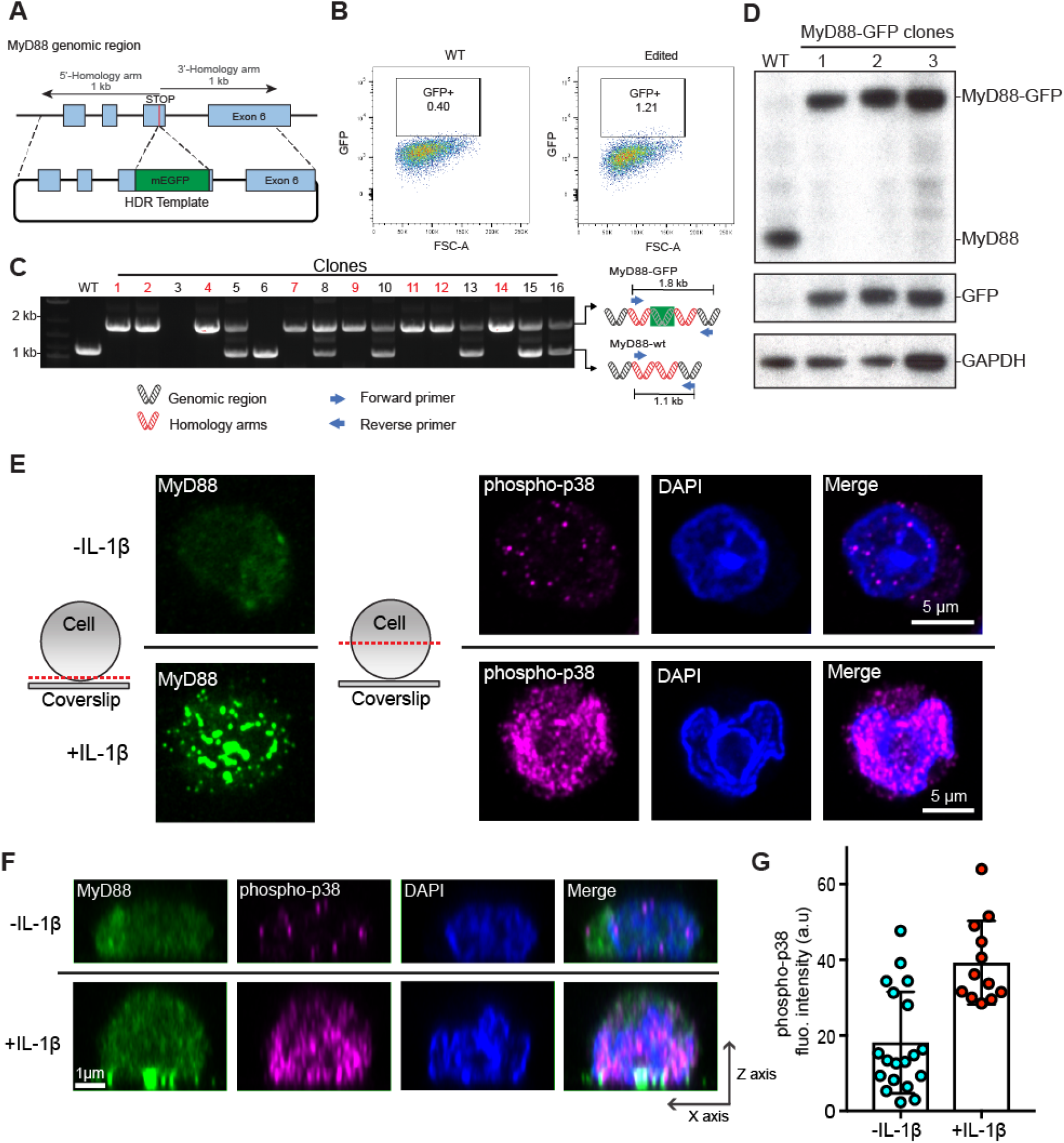
CRISPR/Cas9 gene editing of the MyD88 gene locus with mEGFP and p38 signaling in EL4-MyD88-GFP cells. **A-D) Workflow and validation for CRISPR/Cas9 editing of the MyD88 gene locus with mEGFP.** (A) Schematic of the MyD88 gene locus and homology-directed repair (HDR) template designed to insert a mEGFP open reading frame immediately upstream of the stop codon. (B) FACS results of WT and CRISPR/Cas9-engineered MyD88-GFP EL4 cells. Overlayed box indicates a sorting gate to select EL4-MyD88-GFP cells. (C) PCR screening of CRISPR/Cas9 editing to select homozygous edited cell clones. Schematic shows the primer design for PCR amplification of genomic DNA to detect WT, heterozygous, and homozygous edited EL4 cells. Homozygous MyD88-GFP edited cell clones are labeled in red. Three homozygous clones were retained for Western blot analysis. (D) Western blot analysis of three MyD88-GFP clones. Blots were probed with anti-MyD88, then stripped and reprobed anti-GFP. **E)** To assess the level of MAP kinase pathway activation[, EL4 cells were fixed (60 mins after SLB contact) and stained for MyD88-GFP (green); phospho-p38 (magenta); and DAPI staining of nuclei (blue). Cells were imaged with confocal microscopy. Schematic shows the position of the confocal micrograph slice. Only cells in contact with SLB functionalized with IL1β had increased phospho-p38 nuclear staining intensity. Scale bar, 5 mm. **F)** Reconstructed axial view of cells shown in (E) showing the localization of MyD88 to the cell-SLB contact zone and phospho-p38 staining under IL1β stimulation. Scale bar, 1 mm. **G)** Quantification of phospho-p38 staining intensity. Mean ± SD from n = 12 (with IL1β) and 20 cells (without IL1β).

**Supplementary Figure 2.**
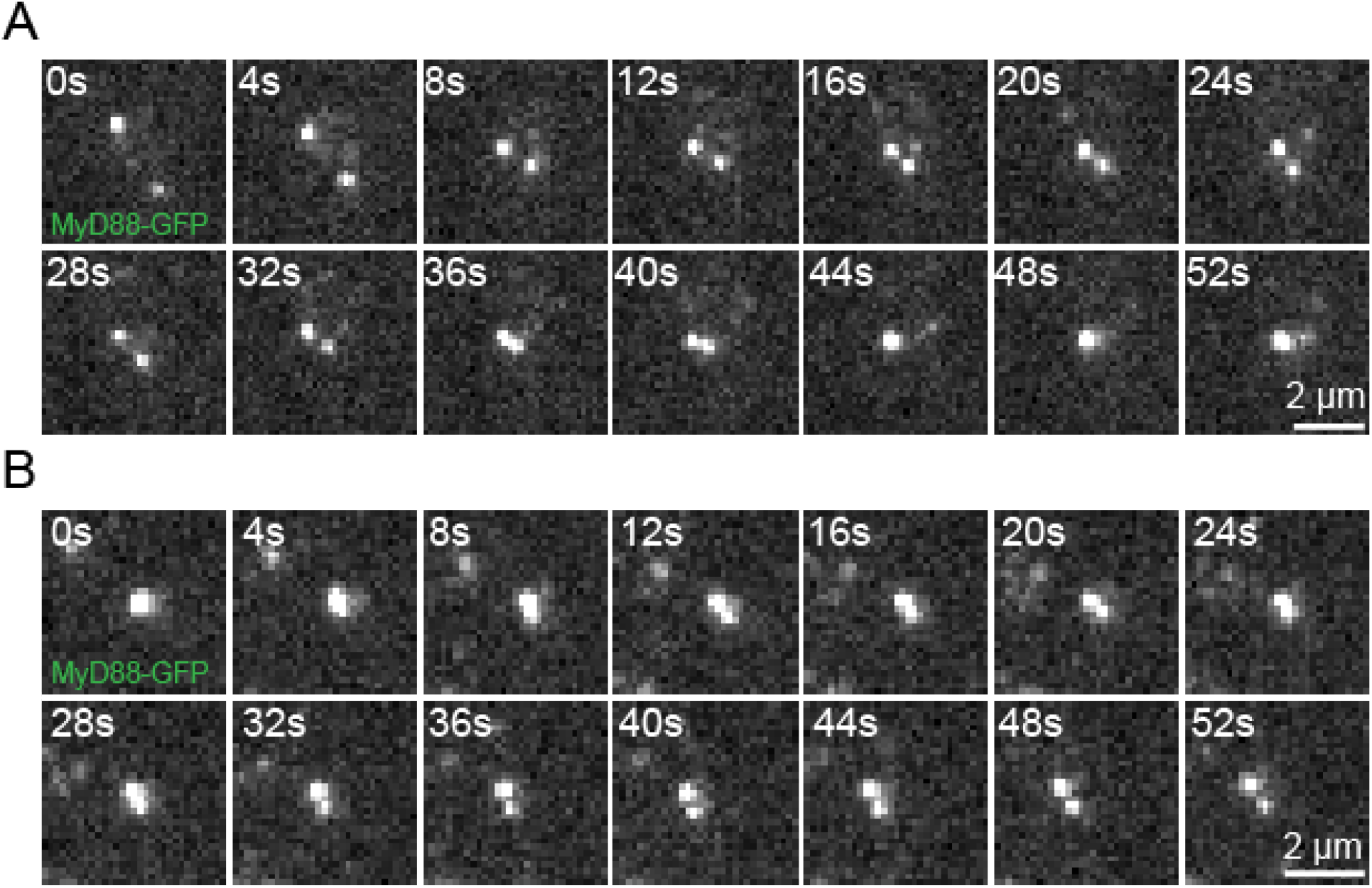
MyD88-GFP puncta can fuse and split. **A)** Example of two MyD88-GFP puncta undergoing fusion**. B)** Example of MyD88-GFP puncta undergoing fission. Scale bar, 2 μm.

**Supplementary Figure 3.**
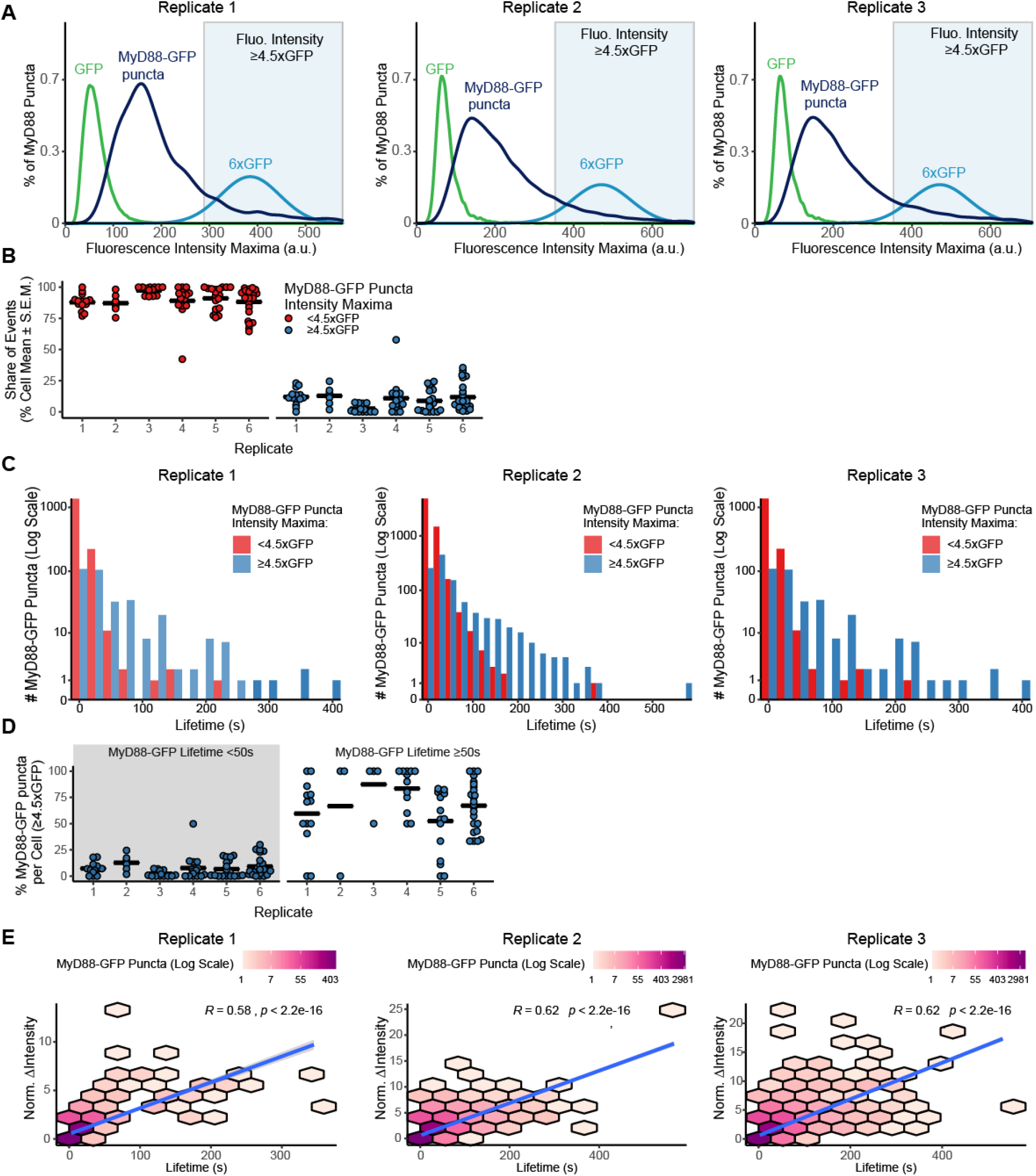
Analysis of MyD88-GFP puncta size, lifetime, and correlation analysis from biological replicates of Fig. 3. **A) Size distribution of MyD88-GFP puncta from three experimental replicates.** Density plot of the maximum fluorescent intensity of MyD88-GFP puncta (dark blue; replicate 1, n = 1,952 puncta from 16 cells; replicate 2, n = 7,637 puncta from 19 cells; replicate 3, n = 11,973 puncta from 24 cells). For comparison, we included the intensity distribution of single GFP fluorophores (green; replicate 1, n = 298,293 GFP particles; replicate 2, n = 7,995 GFP particles; replicate 3, n = 7,995 GFP particles). To estimate the distribution of a 6x GFP multimer (light blue), a Gaussian was fitted to the 1x GFP intensity distribution (see Material and Methods). Blue background shade indicates ≥4.5x GFP. **B) Proportion (%) of the maximum intensity (<4.5x or ≥4.5x GFP) of MyD88-GFP puncta by cell across experimental replicates.** Data points are the proportion of the maximum intensity <4.5x GFP (red) or ≥4.5x GFP (blue) by individual cells from six independent experiments. Percent is the replicate’s proportion of MyD88-GFP puncta with maximum intensity ≥4.5x GFP, n = cell, replicates 1-6: 16% n = 6; 16%, n = 14; 3%, n = 13; 17%, n = 16; 13%, n = 19; 17%, n = 24. **C) Lifetime distribution of MyD88-GFP puncta.** MyD88-GFP lifetime histogram for MyD88-GFP puncta with maximum intensity <4.5x GFP (red) or ≥4.5x GFP (blue; replicate 1, n = 1616 puncta <4.5x GFP and n = 336 ≥4.5x GFP; replicate 2, n = 6553 puncta <4.5x GFP and n = 1084 puncta ≥4.5x GFP; replicate 3, n = 9913 puncta <4.5x GFP and n = 2,060 puncta ≥4.5x GFP). Puncta count is in log scale. **D) Proportion (%) of the lifetime (<50 or <50 s) that are bright MyD88-GFP puncta (≥4.5x GFP) in individual cells across experimental replicates**. Data points are the proportion of individual cells from independent experimental replicates. Percentage of MyD88-GFP puncta with a maximum intensity of ≥4.5x MyD88 that are <50 s (n = cells, from replicate 1 to 6): 7%, n = 14; 13%, n = 6; 2%, n = 13; 8%, n = 16; 7%, n = 19; 9%, n = 24). Percentage of MyD88-GFP puncta with a maximum intensity of ≥4.5x GFP that have lifetimes <50 s (n = cells, replicate 1-6): 59%, n = 14; 67%, n = 6; 88%, n = 13; 83%, n = 16; 52%, n = 19; 67%, n = 24). Long-lived events are more likely to be brighter. Bars in panels F and G represent the replicate mean calculated from the cell means. **E) Correlation between lifetime and intensity growth of MyD88-GFP puncta.** 2D histogram of MyD88-GFP puncta lifetime by change in fluorescent intensity (calculated as maximum intensity minus starting intensity). MyD88-GFP puncta with longer lifetimes have a greater increase in fluorescent intensity. Linear regression line is shown in blue with a 95% confidence interval in gray. There is a statistically-significant strong positive correlation between lifetime and growth (n = puncta, replicate 1-3: R = 0.58, p<0.001 n = 1,952; R = 0.62, p<0.001, n = 7,637; R = 0.62, p<0.001, n = 11,973). Correlations are Spearman’s rank correlation coefficient. Puncta count is shown in log scale.

**Supplementary Figure 4.**
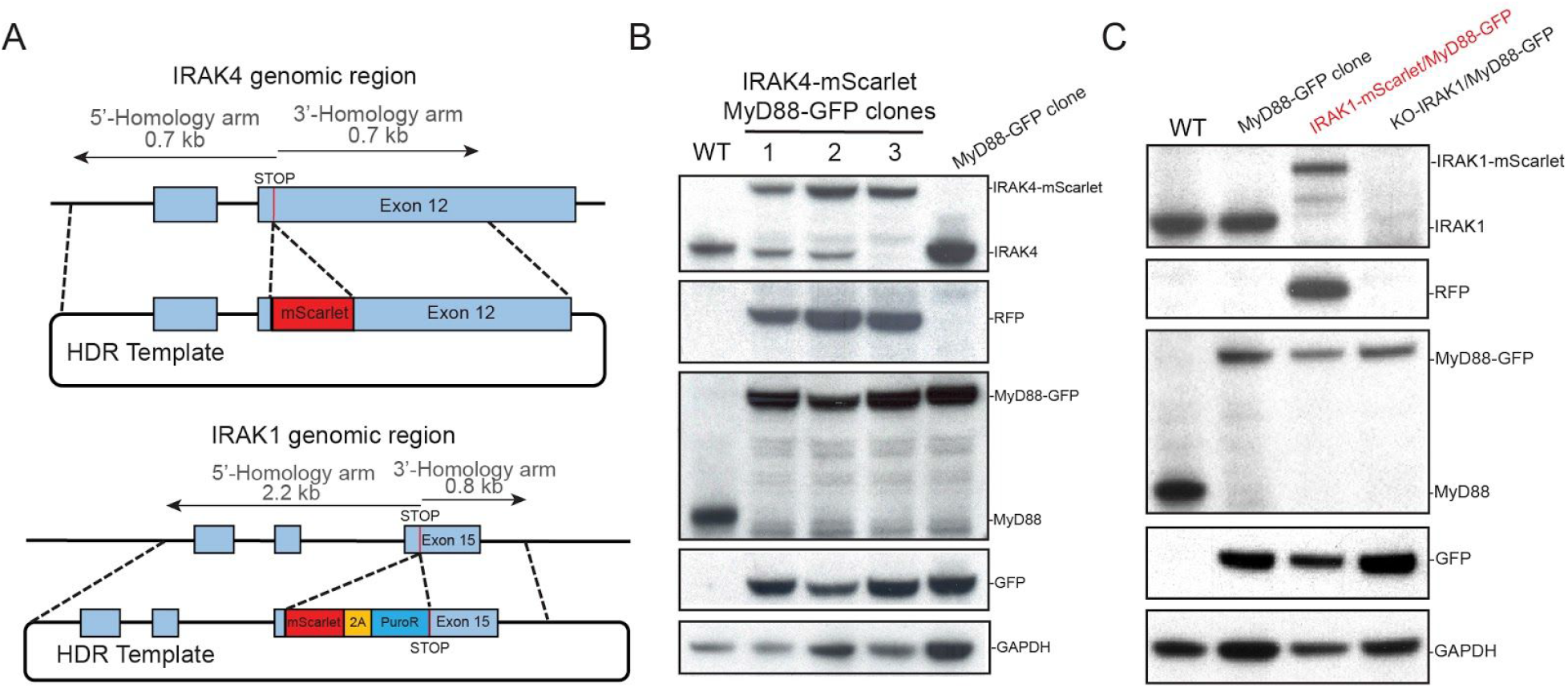
CRISPR/Cas9 gene editing IRAK4 or IRAK1 gene loci with mEGFP and mScarlet. **A)** Schematic of the IRAK4 (top) and IRAK1 (bottom) gene locus and HDR template designed to insert a mScarlet open reading frame immediately upstream of the stop codon. EL4 cells were electroporated with HDR and gRNA/Cas9 plasmids to simultaneously edit MyD88 and IRAK4 or IRAK1 gene loci. Dual edited cells were selected by FACS and PCR (see Fig. S1A-D for workflow, and Materials and Methods). **B)** Western blot analysis of three MyD88-GFP/IRAK4-mScarlet expressing EL4 clones. Lysates were probed with anti-IRAK4, anti-RFP, anti-MyD88, and anti-GFP to confirm editing and insertion of fluorescent protein open reading frames at both gene loci. All data presented in the manuscript were acquired with clone 3. **C)** Western blot analysis of MyD88-GFP/IRAK1-mScarlet expressing EL4 clone. Lysates were probed with anti-IRAK4, anti-RFP, anti-MyD88, and anti-GFP to confirm editing and insertion of mScarlet or mEGFP open reading frames at both gene loci. See supplementary materials for uncropped blots.

**Supplementary Figure 5.**
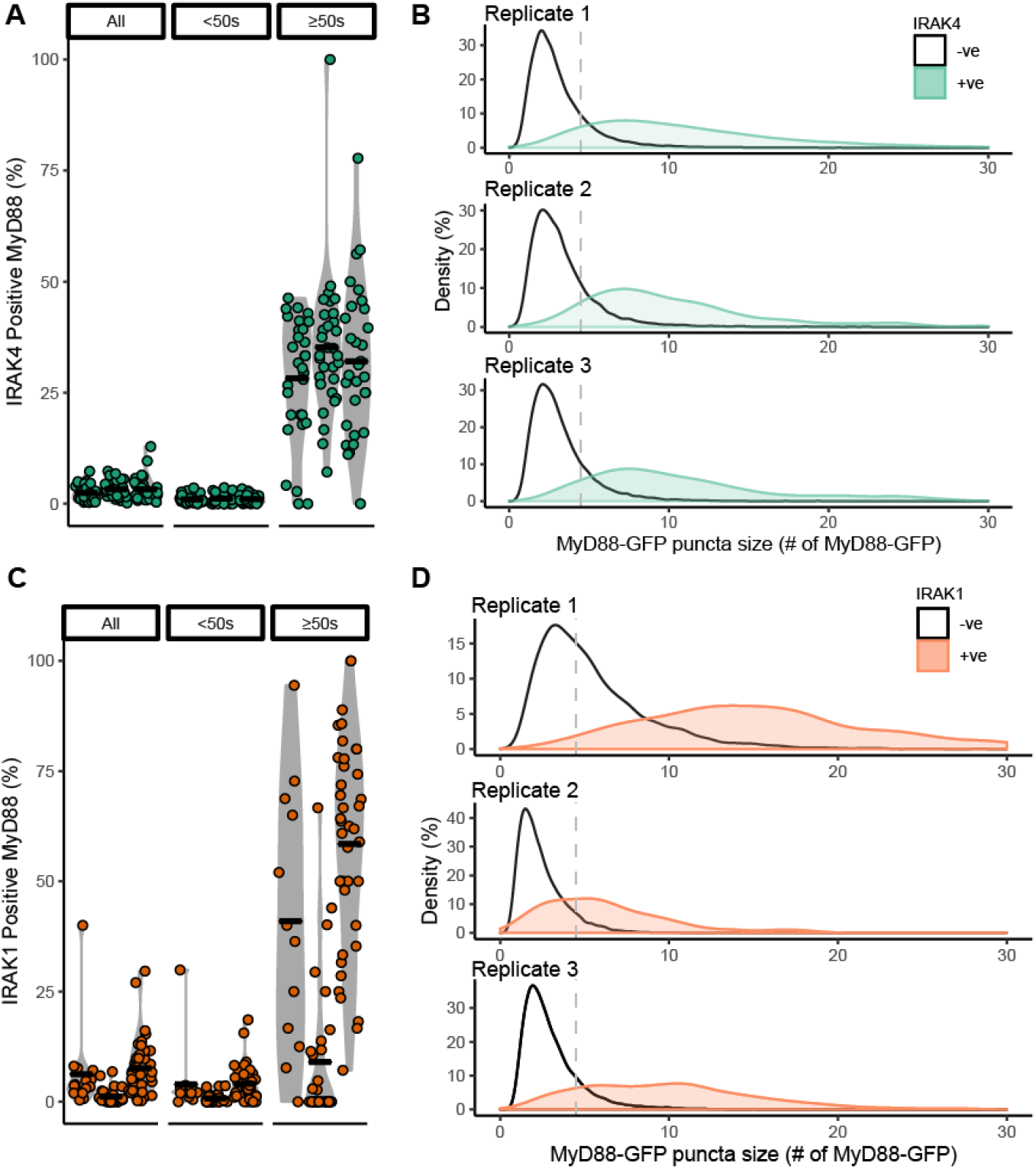
Analysis of MyD88-GFP and IRAK4/1-mScarlet colocalized proportion and puncta size from multiple experimental replicates. Data shown from clonal MyD88-GFP/IRAK4-mScarlet expressing EL4 cells (green, **A-C**) or IRAK1-mScarlet (orange, **D-F**). **(A, C)** Data points are the proportion of individual cells from independent experiments. Bars are experimental replicate means. **(C, F)** Vertical line in (B) and (D) is at 4.5x MyD88-GFP. **A) Percentage (%) of MyD88-GFP puncta that colocalizes with IRAK4-mScarlet, combined, and by MyD88-GFP lifetime (<50 or ≥50 s) per cell across experimental replicates.** Violin plot of the percent of MyD88-GFP puncta that is colocalized with IRAK4-mScarlet, combined, and categorized by lifetime (<50 or ≥50 s). Few tracks recruit IRAK4 (“All”, n = cells, replicates 1-3: 2.4%, n = 30; 3.2%, n = 31; 3.3%, n = 30). It is especially evident in MyD88-GFP puncta that persist for <50 s (“<50 s”, n = cells, replicates 1-3: 1.0%, n = 30; 1.2%, n = 31; 1.1%, n = 30). However, MyD88-GFP puncta that persist for ≥50 s are more likely to recruit IRAK4 (“≥50 s”, n = cells, replicates 1-3: 28%, n = 30; 35%, n = 31; 32%, n = 30). **B) Maximum MyD88-GFP size of IRAK4-mScarlet colocalized and non-colocalized puncta across experimental replicates.** Density plot of MyD88-GFP size categorized as colocalized (blue) or non-colocalized (red) with IRAK4-GFP. MyD88-GFP puncta colocalized with IRAK4-mScarlet are brighter (mean colocalized vs not colocalized, n = puncta, replicates 1-3: 11x MyD88-GFP, n = 835 vs. 3.1x MyD88-GFP, n = 29,072; 10x MyD88-GFP, n = 601 vs. 3.3x MyD88-GFP, n = 17,221; 11.7x MyD88-GFP, n = 552 vs. 3.3x MyD88-GFP, n = 17,854). **C) Percentage (%) of MyD88-GFP puncta that colocalizes with IRAK1-mScarlet, combined and by MyD88-GFP lifetime (<50 or ≥50 s) per cell across experimental replicates.** Few tracks recruit IRAK1 (“All”, n = cells, replicates 1-3: 6.3%, n = 15; 1.5%, n = 32; 7.6%, n = 40). It is especially evident in MyD88-GFP puncta that persist for ≥50 s (“≥50 s”, n = cells, replicates 1-3: 3.9%, n = 15; 0.74%, n = 32; 4.1%, n = 40). However, MyD88 puncta that persist for <50 s are more likely to recruit IRAK1 (“<50 s”, n = cells, replicates 1-3: 41%, n = 15; 9.0%, n = 32; 59%, n = 40). **D) Maximum MyD88-GFP size of IRAK1-mScarlet colocalized and non-colocalized puncta across experimental replicates.** Density plot of MyD88-GFP size categorized as colocalized (blue) or not colocalized (red) with IRAK1-GFP. MyD88-GFP puncta colocalized with IRAK1-mScarlet are brighter (mean colocalized vs not colocalized, n = puncta, replicates 1-3: 19x MyD88-GFP, n = 291 vs. 5.6x MyD88-GFP, n = 5,401; 6.2x MyD88-GFP, n = 314 vs. 2.4x MyD88-GFP, n = 14,125; 10xMyD88-GFP, n = 1,439 vs. 3.1x MyD88-GFP, n = 16,834).

**Supplementary Figure 6.**
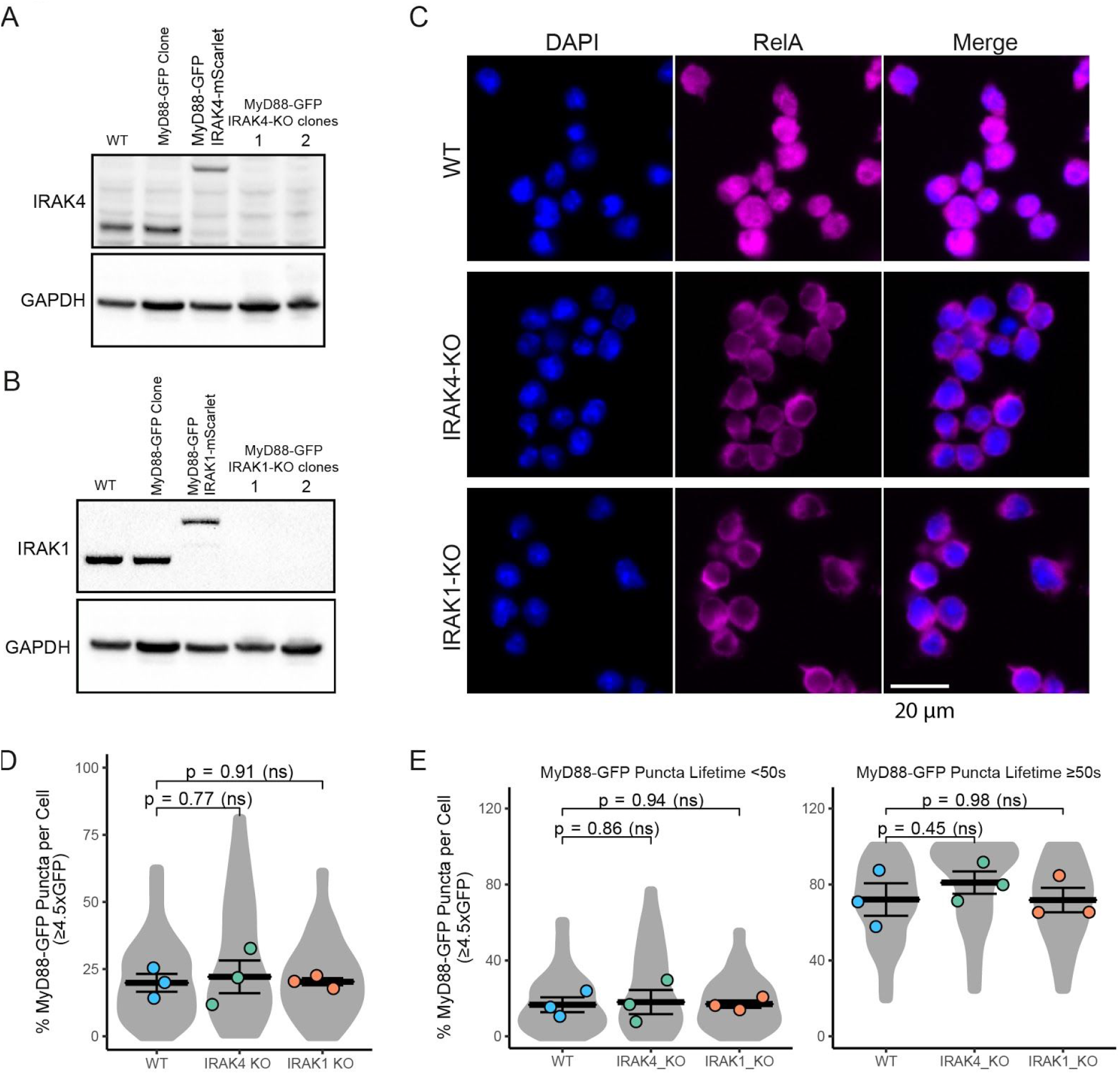
CRISPR/Cas9 KO IRAK4 and IRAK1 EL4 cell lines, and MyD88-GFP dynamics in KO cell lines. **A-B) Validation of IRAK4 and IRAK1 KO cell lines.** Western blot analysis of two monoclonal EL4 IRAK4 KO (A) and IRAK1 KO (B) cell lines. **C) RelA does not translocate to the nucleus in IRAK4 KO and IRAK1 KO cell lines when stimulated with IL1β.** EL4 cell lines (30 mins after addition to IL1β-labeled SLBs) were fixed and stained for RelA (magenta); DAPI stain nuclei (blue). Scale bar, 20 μm. **D) The percentage per cell of large MyD88 oligomers in WT, IRAK4 KO and IRAK1 KO EL4 cell lines.** Quantification of the percentage (%) per cell of MyD88-GFP puncta with a maximum intensity ≥ 4.5x GFP. Violin plots show the distribution of the cell data. Data points superimposed on violin plots are the averages of independent experiments. Bars represent mean ± SEM (n = 3 experimental replicates, with > 10 cells measured per replicate). **E) The percentage per cell of large MyD88 oligomers is equivalent for short- and long-lived MyD88-GFP puncta across WT and KO cell lines.** Quantification of the proportion (%) per cell of MyD88-GFP puncta with a maximum intensity of ≥4.5x GFP categorized by lifetimes of < or ≥ 50 s. Violin plots show the distribution of the cell data. Data points superimposed on the violin plots are the averages from independent experiments. Bars represent mean ± SEM (n = 3 experimental replicates, with > 10 cells measured per replicates). WT and KO image means were compared using an unpaired Student’s t-test.

**Supplementary Figure 7.**
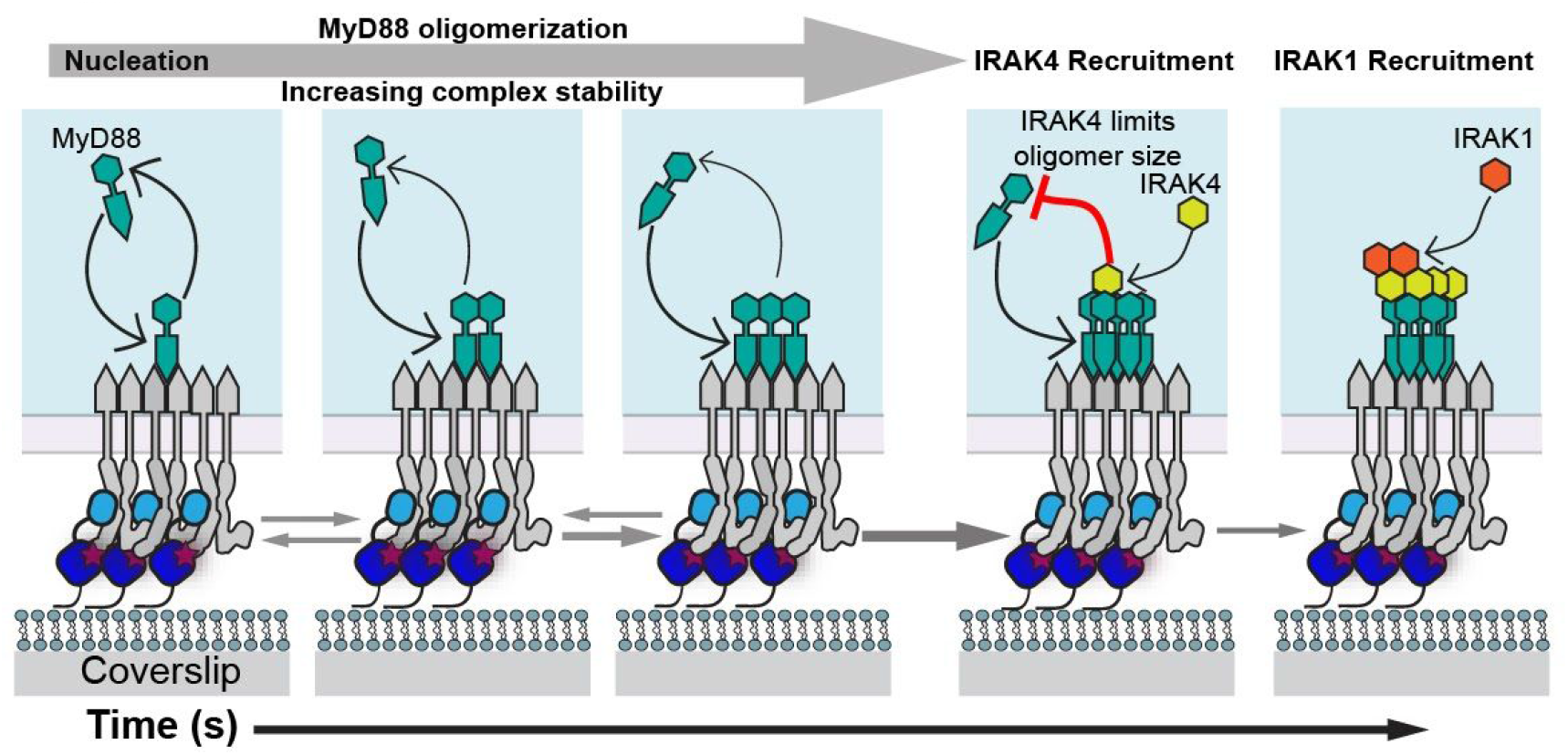
Model describing the nucleation and assembly of Myddosomes. MyD88 oligomer stability and size regulates the incorporation of IRAK4 and IRAK1 in Myddosomes. MyD88 recruitment to the IL1-bound IL1R nucleates MyD88 oligomerization. Initially, the small oligomer of MyD88 is unstable and can disassemble into the cytosol. However, as MyD88 oligomer size increases, so does complex stability, and the formation of larger MyD88 complexes triggers downstream signalling in the form of the sequential recruitment of IRAK4 followed by IRAK1. Critically, IRAK4 limits MyD88 oligomer size, and inhibits further MyD88 oligomerization.

## References

Akira S, Uematsu S, Takeuchi O. 2006. Pathogen recognition and innate immunity. Cell 124:783–801.

Bindels DS, Haarbosch L, van Weeren L, Postma M, Wiese KE, Mastop M, Aumonier S, Gotthard G, Royant A, Hink MA, Gadella TWJ. 2017. mScarlet: a bright monomeric red fluorescent protein for cellular imaging. Nat Methods 14:53–56.

Bird TA, Gearing AJ, Saklatvala J. 1988. Murine interleukin 1 receptor. Direct identification by ligand blotting and purification to homogeneity of an interleukin 1-binding glycoprotein. J Biol Chem 263:12063–12069.

Biswas KH, Groves JT. 2019. Hybrid Live Cell-Supported Membrane Interfaces for Signaling Studies. Annu Rev Biophys 48:537–562.

Bonham KS, Orzalli MH, Hayashi K, Wolf AI, Glanemann C, Weninger W, Iwasaki A, Knipe DM, Kagan JC. 2014. A promiscuous lipid-binding protein diversifies the subcellular sites of toll-like receptor signal transduction. Cell 156:705–716.

DeFelice MM, Clark HR, Hughey JJ, Maayan I, Kudo T, Gutschow MV, Covert MW, Regot S. 2019. NF-κB signaling dynamics is controlled by a dose-sensing autoregulatory loop. Sci Signal 12:eaau3568.

Deng L, Wang C, Spencer E, Yang L, Braun A, You J, Slaughter C, Pickart C, Chen ZJ. 2000. Activation of the IkappaB kinase complex by TRAF6 requires a dimeric *ubiquitin-conjugating enzyme complex and a unique polyubiquitin chain*. Cell 103:351–361.

Ferrao R, Wu H. 2012. Helical assembly in the death domain (DD) superfamily. Curr Opin Struct Biol 22:241–247.

Fu T-M, Li Y, Lu A, Li Z, Vajjhala PR, Cruz AC, Srivastava DB, DiMaio F, Penczek PA, Siegel RM, Stacey KJ, Egelman EH, Wu H. 2016. Cryo-EM Structure of Caspase-8 Tandem DED Filament Reveals Assembly and Regulation Mechanisms of the Death-Inducing Signaling Complex. Mol Cell 64:236–250.

Gay NJ, Symmons MF, Gangloff M, Bryant CE. 2014. Assembly and localization of Toll-like receptor signalling complexes. Nature Publishing Group 14:546–558.

Hardiman G, Rock FL, Balasubramanian S, Kastelein RA, Bazan JF. 1996. Molecular characterization and modular analysis of human MyD88. Oncogene 13:2467–2475.

Horng T, Barton GM, Flavell RA, Medzhitov R. 2002. The adaptor molecule TIRAP provides signalling specificity for Toll-like receptors. Nature 420:329–333.

Jacobs SA, Diem MD, Luo J, Teplyakov A, Obmolova G, Malia T, Gilliland GL, O’Neil KT. 2012. Design of novel FN3 domains with high stability by a consensus sequence approach. Protein Eng Des Sel 25:107–117.

Kagan JC, Magupalli VG, Wu H. 2014. SMOCs: supramolecular organizing centres that control innate immunity. Nature Publishing Group 14:821–826.

Kagan JC, Medzhitov R. 2006. Phosphoinositide-mediated adaptor recruitment controls Toll-like receptor signaling. Cell 125:943–955.

Kang M, Andreani M, Kenworthy AK. 2015. Validation of Normalizations, Scaling, and Photofading Corrections for FRAP Data Analysis. PLoS One 10:e0127966.

Kaplanski G, Farnarier C, Kaplanski S, Porat R, Shapiro L, Bongrand P, Dinarello CA. 1994. Interleukin-1 induces interleukin-8 secretion from endothelial cells by a juxtacrine mechanism. Blood 84:4242–4248.

Kawagoe T, Sato S, Matsushita K, Kato H, Matsui K, Kumagai Y, Saitoh T, Kawai T, Takeuchi O, Akira S. 2008. Sequential control of Toll-like receptor-dependent responses by IRAK1 and IRAK2. Nat Immunol 9:684–691.

Keeble AH, Banerjee A, Ferla MP, Reddington SC, Anuar INAK, Howarth M. 2017. Evolving Accelerated Amidation by SpyTag/SpyCatcher to Analyze Membrane Dynamics. Angew Chem Int Ed Engl 56:16521–16525.

Latty SL, Sakai J, Hopkins L, Verstak B, Paramo T, Berglund NA, Cammorota E, Cicuta P, Gay NJ, Bond PJ, Klenerman D, Bryant CE. 2018. Activation of Toll-like receptors nucleates assembly of the MyDDosome signaling hub. Elife 7.

Latz E, Visintin A, Lien E, Fitzgerald KA, Monks BG, Kurt-Jones EA, Golenbock DT, Espevik T. 2002. Lipopolysaccharide rapidly traffics to and from the Golgi apparatus with the toll-like receptor 4-MD-2-CD14 complex in a process that is distinct from the initiation of signal transduction. J Biol Chem 277:47834–47843.

Legland D, Arganda-Carreras I, Andrey P. 2016. MorphoLibJ: integrated library and plugins for mathematical morphology with ImageJ. Bioinformatics 32:3532–3534.

Lin S-C, Lo Y-C, Wu H. 2010. Helical assembly in the MyD88-IRAK4-IRAK2 complex in TLR/IL-1R signalling. Nature 465:885–890.

Lu A, Magupalli VG, Ruan J, Yin Q, Atianand MK, Vos MR, Schröder GF, Fitzgerald KA, Wu H, Egelman EH. 2014. Unified polymerization mechanism for the assembly of ASC-dependent inflammasomes. Cell 156:1193–1206.

Medzhitov R, Janeway C Jr. 2000. Innate immune recognition: mechanisms and pathways. Immunol Rev 173:89–97.

Medzhitov R, Preston-Hurlburt P, Kopp E, Stadlen A, Chen C, Ghosh S, Janeway CA. 1998. MyD88 is an adaptor protein in the hToll/IL-1 receptor family signaling pathways. Mol Cell 2:253–258.

Mohapatra L, Goode BL, Jelenkovic P, Phillips R, Kondev J. 2016. Design Principles of Length Control of Cytoskeletal Structures. Annu Rev Biophys 45:85–116.

Moncrieffe MC, Bollschweiler D, Li B, Penczek PA, Hopkins L, Bryant CE, Klenerman D, Gay NJ. 2020. MyD88 Death-Domain Oligomerization Determines Myddosome Architecture: Implications for Toll-like Receptor Signaling. Structure.

Motshwene PG, Moncrieffe MC, Grossmann JG, Kao C, Ayaluru M, Sandercock AM, Robinson CV, Latz E, Gay NJ. 2009. An oligomeric signaling platform formed by *the Toll-like receptor signal transducers MyD88 and IRAK-4*. J Biol Chem 284:25404–25411.

Ngo VN, Young RM, Schmitz R, Jhavar S, Xiao W, Lim K-H, Kohlhammer H, Xu W, Yang Y, Zhao H, Shaffer AL, Romesser P, Wright G, Powell J, Rosenwald A, Muller-Hermelink HK, Ott G, Gascoyne RD, Connors JM, Rimsza LM, Campo E, Jaffe ES, Delabie J, Smeland EB, Fisher RI, Braziel RM, Tubbs RR, Cook JR, Weisenburger DD, Chan WC, Staudt LM. 2011. Oncogenically active MYD88 mutations in human lymphoma. Nature 470:115–119.

O’Carroll A, Chauvin B, Brown JWP, Meagher A, Coyle J, Schill J, Bhumkhar A, Hunter DJB, Ve T, Kobe B, Sierecki E, Gambin Y. 2018. Pathological mutations differentially affect the self-assembly and polymerisation of the innate immune system signalling adaptor molecule MyD88. BMC Biol 16:149.

O’Neill LA, Bird TA, Saklatvala J. 1990. How does interleukin 1 activate cells? Interleukin 1 signal transduction. Immunol Today 11:392–394.

O’Neill LAJ. 2008. The interleukin-1 receptor/Toll-like receptor superfamily: 10 years of progress. Immunol Rev 226:10–18.

O’Neill LAJ, Bowie AG. 2007. The family of five: TIR-domain-containing adaptors in Toll-like receptor signalling. Nat Rev Immunol 7:353–364.

Park HH, Logette E, Raunser S, Cuenin S, Walz T, Tschopp J, Wu H. 2007. Death domain assembly mechanism revealed by crystal structure of the oligomeric PIDDosome core complex. Cell 128:533–546.

Suzuki N, Suzuki S, Duncan GS, Millar DG, Wada T, Mirtsos C, Takada H, Wakeham A, Itie A, Li S, Penninger JM, Wesche H, Ohashi PS, Mak TW, Yeh W-C. 2002. Severe impairment of interleukin-1 and Toll-like receptor signalling in mice lacking IRAK-4. Nature 416:750–756.

Tan Y, Kagan JC. 2019. Innate Immune Signaling Organelles Display Natural and Programmable Signaling Flexibility. Cell 177:384–398.e11.

Taylor MJ, Husain K, Gartner ZJ, Mayor S, Vale RD. 2017. A DNA-Based T Cell *Receptor Reveals a Role for Receptor Clustering in Ligand Discrimination*. Cell 169:108–119.e20.

Tinevez J-Y, Perry N, Schindelin J, Hoopes GM, Reynolds GD, Laplantine E, Bednarek SY, Shorte SL, Eliceiri KW. 2017. TrackMate: An open and extensible platform for single-particle tracking. Methods 115:80–90.

Vayttaden SJ, Smelkinson M, Ernst O, Carlson RJ, Sun J, Bradfield C, Dorrington MG, Liang J, Bouladoux N, Gottschalk RA, Oh K-S, Pegoraro G, Ganesan S, De Nardo D, Latz E, Belkaid Y, Varma RR, Fraser IDC. 2019. IRAK1-mediated coincidence *detection of microbial signals licenses inflammasome activation*. bioRxiv. doi: 10.1101/2019.12.26.888776

Ve T, Vajjhala PR, Hedger A, Croll T, DiMaio F, Horsefield S, Yu X, Lavrencic P, Hassan Z, Morgan GP, Mansell A, Mobli M, O’Carroll A, Chauvin B, Gambin Y, Sierecki E, Landsberg MJ, Stacey KJ, Egelman EH, Kobe B. 2017. Structural basis of TIR-domain-assembly formation in MAL- and MyD88-dependent TLR4 signaling. Nat Struct Mol Biol 24:743–751.

Wang L, Yang JK, Kabaleeswaran V, Rice AJ, Cruz AC, Park AY, Yin Q, Damko E, Jang SB, Raunser S, Robinson CV, Siegel RM, Walz T, Wu H. 2010. The Fas-FADD death domain complex structure reveals the basis of DISC assembly and disease mutations. Nat Struct Mol Biol 17:1324–1329.

Wu H. 2013. Higher-order assemblies in a new paradigm of signal transduction. Cell 153:287–292.

