## Supplemental Materials for "On demand MyD88 oligomerization is controlled by IRAK4 during Myddosome signaling"

+ Equal author contribution

### Table S1. Key resources table

| REAGENT or RESOURCE | SOURCE | IDENTIFIER |
|---|---|---|
| **Antibodies** | | |
| Goat polyclonal anti-MyD88 | R and D Systems | Cat# AF3109; RRID:AB_2146703 |
| Rabbit polyclonal anti-IRAK4 | Cell Signaling Technology | Cat# 4363; RRID:AB_2126429 |
| Rabbit monoclonal anti-IRAK1 (D51G7) | Cell Signaling Technology | Cat# 4504; RRID:AB_1904032 |
| Rabbit polyclonal anti-GFP | ChromoTek | Cat# PABG1; RRID:AB_2749857 |
| Mouse monoclonal anti-RFP | ChromoTek | Cat# 6g6; RRID:AB_2631395 |
| Rabbit polyclonal anti-GAPDH | AbFrontier | Cat# LF-PA0018; RRID:AB_1616734 |
| Rabbit monoclonal anti-NF-κB p65 | Cell Signaling Technology | Cat# 8242; RRID: AB_10859369 |
| Rabbit monoclonal anti-Phospho-p38 MAPK | Cell Signaling Technology | Cat# 4511; RRID: AB_2139682 |
| Camelid sdAb FluoTag®-X4 anti-GFP-Atto488 | NanoTag Biotechnologies GmbH | Cat# N0304-At488; |
| **Bacterial and Virus Strains** | | |
| BL21 Rosetta | Novagen | Cat# 70954 |
| NEB Stable | NEB | Cat# C3040 |
| **Chemicals, Peptides, and Recombinant Proteins** | | |
| His10-mScarlet-i-Tencon-SpycatcherV2-IL1β | This study | |
| His10-Halo-Tencon-SpycatcherV2-IL1β | This study | |
| Puromycin dihydrochloride | Thermo | Cat# A1113803 |
| Blasticidin | InvivoGen | Cat# ant-bl-05 |

| | | |
|---|---|---|
| JF646-HaloTag ligand | Luke Lavis, HHMI | |
| **Experimental Models: Cell Lines** | | |
| Mouse: EL4.NOB-1 | ECACC | 87020408 |
| Mouse: EL4 MyD88-GFP | This paper | |
| Mouse: EL4 IRAK1-mScarlet/MyD88-GFP | This paper | |
| Mouse: EL4 IRAK4-mScarlet/MyD88-GFP | This paper | |
| Mouse: EL4 IRAK1-KO/MyD88-GFP clone 1 | This paper | |
| Mouse: EL4 IRAK1-KO/MyD88-GFP clone 2 | This paper | |
| Mouse: EL4 IRAK4-KO/MyD88-GFP clone 1 | This paper | |
| Mouse: EL4 IRAK4-KO/MyD88-GFP clone 2 | This paper | |
| **sgRNA sequences** | | |
| MyD88 GFP knock in gRNA1: CCTGCCCTGAAGATGACCCT | This study | |
| MyD88 GFP knock in gRNA2: TGACATACCTAGGGCTCCCA | This study | |
| IRAK1 mScarlet-i knock in gRNA: TTAGAACCTGAAAAGAGCCA | This study | |
| IRAK4 mScarlet-i knock in gRNA: TTCAAGACATCGGCTTAACC | This study | |
| IRAK1 knockout gRNA: GCCTGGAACCACAGGCtcccc | This study | |
| IRAK4 knockout gRNA: GAGGTTGCGTATGTATGTCGA | This study | |
| **Recombinant DNA** | | |
| pMK | Thermo | GeneArt vector |
| pX330-U6-Chimeric_BB-CBh-hSpCas9 | Addgene | Plasmid #42230 |
| pX459V2.0-HypaCas9 | Addgene | Plasmid #108294 |
| pMK-MyD88-GFP-HDRtemp | This paper | |

| | | |
|---|---|---|
| pX330-MyD88-gRNA-1 | This paper | |
| pX330-MyD88-gRNA-2 | This paper | |
| pMK-IRAK1-mScarlet-P2A-Puro-HDRtemp | This paper | |
| pX330-IRAK1-gRNA | This paper | |
| pMK-IRAK4-mScarlet-HDRtemp | This paper | |
| pX330-IRAK4-gRNA | This paper | |
| pMK-IRAK1-KO-BlastR-Stop-HDRtemp | This paper | |
| pX459-IRAK1-KO-gRNA | This paper | |
| pMK-IRAK4-KO-BlastR-Stop-HDRtemp | This paper | |
| pX459-IRAK4-KO-gRNA | This paper | |
| **Software and Algorithms** | | |
| Fiji | NIH | https://fiji.sc/ |
| MATLAB | MathWorks | https://www.mathworks.com/ |
| R | CRAN | https://cran.r-project.org/ |
| ggplot2 | tidyverse | https://ggplot2.tidyverse.org/ |
| Fiji/MATLAB/R scripts from image analysis | This study | https://gitlab.com/Marcus_Taylor/myddosome-dynamics-pipeline |

### Table S2. PCR primers for *MyD88* sequence validation

| Full Primer Name | 5'-3' Sequence | Amplified Region | |
|---|---|---|---|
| | | WT-*MyD88*: | *MyD88*-GFP: |
| MyD88-seq $F_{in}$ | CTGGACCCGCCTTGCCAAGGC | 1.1kb | 1.8kb |

| | | | |
|---|---|---|---|
| MyD88-seq $R_{out}$ | GAAGATGCAAACCTCGCTGC TGGGG | | |
| mGFP-seq F | TGCCCGACAACCACTACCTG AGC | - | 1.2kb |
| MyD88-seq $R_{out}$ | GAAGATGCAAACCTCGCTGC TGGGG | | |

### Table S3. PCR primers for *IRAK4* sequence validation

| Full Primer Name | 5'-3' Sequence | Amplified Region | |
|---|---|---|---|
| | | **WT-*IRAK4:*** | ***IRAK4-mScarlet*:** |
| IRAK4-seq $F_{out}$ | GAGCCACCTTTCAACCTTCC GT | - | 1.0kb |
| mScarlet-seq R | CACCTTCAGCTTGGCGGTCT G | | |
| IRAK4-seq $F_{out}$ | GAGCCACCTTTCAACCTTCC GT | 0.9kb | 1.6kb |
| IRAK4-seq $R_{in}$ | GCACTATGCTACCATGTTAA ACATAAAGCGC | | |

### Table S4. PCR primers for *IRAK1* sequence validation

| Full Primer Name | 5'-3' Sequence | Amplified Region | |
|---|---|---|---|
| | | **WT-*IRAK1:*** | ***IRAK1-mScarlet*:** |
| mScarlet-seq F | GACCGCAAGTTGGACATCAC CT | - | 1.7kb |
| IRAK1-seq $R_{out}$ | AGCCTGCCTAGGCAGGCAG GTAGTC | | |
| IRAK1-seq $F_{in}$ | CAAAGTTCTGTGCTCATGGT TCATGTCAGGG | 1.0kb | 2.5kb |
| IRAK1-seq $R_{out}$ | AGCCTGCCTAGGCAGGCAG GTAGTC | | |

**Table S5. PCR primers for *IRAK4* KO sequence validation**

| Full Primer Name | 5'-3' Sequence | Amplified Region | |
|---|---|---|---|
| | | WT-*IRAK4:* | *IRAK4-BlastR-KO* |
| IRAK4-KO-seq $F_{in}$ | TGGGTCGGAGTGAAAGCTGCTC | - | 0.67kb |
| BlastR-seq R | GCTGTCCATCACTGTCCTTCACTATG | | |
| IRAK4-KO-seq $F_{in}$ | TGGGTCGGAGTGAAAGCTGCTC | 1.1kb | 1.5kb |
| IRAK4-KO-seq $R_{out}$ | GACACTTGCTGGAAGGTCAATATGG | | |

**Table S6. PCR primers for *IRAK1* KO sequence validation**

| Full Primer Name | 5'-3' Sequence | Amplified Region | |
|---|---|---|---|
| | | WT-*IRAK1:* | *IRAK1-BlastR-KO* |
| BlastR-seq F | GGACAGTGATGGACAGCCGAC | - | 1.4kb |
| IRAK1-KO-seq $R_{out\_v1}$ | CTTCCAGCAGTCAAGCCCAGAGA | | |
| IRAK1-KO-seq $F_{in}$ | AGGCCGCGGAGGGCAAGATG | 1.0kb | 1.4kb |
| IRAK1-KO-seq $R_{out\_v2}$ | GGAAACAGGGAGTGGAACCTGGA | | |

**Table S7. Sequences of the *HDR templates***

HDR arms; mEGFP ORF, mScarlet-i ORF, 2A-PuroR ORF, BlastR ORF, STOP Codon, START Codon.

| MyD88-mEGFP HDR template |
|---|
| atGCCTCCATCATAGTTAACCGGGATTTCATCTGGGAGGAAGTTATCTGTTCACTGGTAGA |

GAGGGCATGTATATGACATTGCTTTGATATGGATACAGGCCCAGGTTCCCTTGATGGAAG
ACTCCAGGTTGGGCTCCTTCCAGCCTTCTGCAGAGGCTGATTGATTCCCTTGTCCCCTGT
CCTCAGGACAAACGCCGGAACTTTTCGATGCCTTTATCTGCTACTGCCCCAACGATATCG
AGTTTGTGCAGGAGATGATCCGGCAACTAGAACAGACAGACTATCGGCTTAAGTTGTGT
GTGTCCGACCGTGACGTCCTGCCGGGCACCTGTGTCTGGTCCATTGCCAGCGAGCTAAT
TGAGAAAAGGTTGGTTAAACATCTAAGAGGGTAGGTGGGTGAATGCATGAAACCCAGAG
GTCCAGATGCAAGGACTGTCCTGCTAGCTGGGCTCTGTCCCGCCTGGGTAATGTAGTCC
TTCCTGACCCCATCCTCTGAAGGAAGTCACCGCAGTGCCACTCTCCCTCAGGTGTCGCC
GCATGGTGGTGGTTGTTTCTGACGATTATCTACAGAGCAAGGAATGTGACTTCCAGACCA
AGTTTGCACTCAGCCTGTCTCCAGGTAAGCTTAGGCCTGCTTTGGTCAAAGAGAGAGTA
GAGATATAGCCTTAGGATGATAGTCCAGGAATGCAAAACCAAAGCACTACAGATCTCTGA
GGACGGACCTGTGTACTTCCTTATGTAGTGGATATGTATCATGGATACCTGGTTGGTGGA
GAGCTGGTGTTAGCCCGTGCTTTAGGGACTGAGCCTGTCCCACCCTAGGGCCCCACGT
GGTCCTAATACCACACCCTTTGGCCTTCAGGTGTCCAACAGAAGCGACTGATTCCTATTA
AATACAAGGCGATGAAGAAGGACTTTCCCAGTATCCTGCGGTTCATCACTATATGCGACT
ATACCAACCCTTGCACCAAGTCCTGGTTCTGGACCCGCCTTGCCAAGGCTTTGTCCCTG
CCCGGAGGATCTGGTGGATCAGGTGGAAGTgtgagcaagggcgaggagctgttcaccggggtggtgccca
tcctggtcgagctggacggcgacgtaaacggccacaagttcagcgtgtccggcgagggcgagggcgatgccacctacggcaa
gctgaccctgaagttcatctgcaccaccggcaagctgcccgtgccctggcccaccctcgtgaccaccctgacctacggcgtgcag
tgcttcagccgctaccccgaccacatgaagcagcacgacttcttcaagtccgccatgcccgaaggctacgtccaggagcgcacc
atcttcttcaaggacgacggcaactacaagacccgcgccgaggtgaagttcgagggcgacaccctggtgaaccgcatcgagct
gaagggcatcgacttcaaggaggacggcaacatcctggggcacaagctggagtacaactacaacagccacaacgtctatatca
tggccgacaagcagaagaacggcatcaaggtgaacttcaagatccgccacaacatcgaggacggcagcgtgcagctcgccg
accactaccagcagaacacccccatcggcgacggccccgtgctgctgcccgacaaccactacctgagcacccagtccaagctg
agcaaagaccccaacgagaagcgcgatcacatggtcctgctggagttcgtgaccgccgccgggatcactctcggcatggacga
gctgtacaagtgaAGATGACaCTGaGAACCCTAtGTATGTCAGTCTGTCTGTGTTCTTCCGCTTG
CCTCCTTTGACACTGTAGTGGGGAGCTTGTGGTCCGTCTATCCCTAGACATCTACAGTAG
CCAGATGTCATCTCTCAAGTCCTTACGGAACCAGTGACTGCAAGTGACATTATGACTTGC
CGGGTTGCCAGCCAGGACAGTGTTTATCAGGGCTGCATGAGTCTAAGCGAAGGACTAGT
TGAGCTACATCTCAGACATCTTTGGCCTCAGGGTCTCCTCCTGAGAAACAGGATATGGA
GATCAGGCAGGGAAATAGTGAGGCACTCTTCTGGCTATCCTAGGAGACCCAGTGTGGGC
TGAAGGGGAGCTGAAGGAGCACACACTGGCTGTTAAGGAGCACCACAGGCTTGCCAGG
ACAGCTGCTTTGACCTGCCAACCTTAAACTCCCTTTCTTCCTCCCACAGGGCAGAGGGG
AAGATGAGACTGATGCGGAGCCAGATTCTCTGATGCCGTCCTGTCTACATCTTTGACTCC
CCTGGGCTCAACCCGTGTTCAATGATGACTGGCCTGAGCAACTAGGACTGCCTTTCCTC
CCAGCCACCCATGCCTGTGCACGCACCTCAGTACACACATGCCTCCTCGCACACACAGG
CATCTGCATATGTGTGTTTCCTTTGGGACAGCTCCCAAGGATAGCTGAGTGGAAGAGTTC
TATCATCAAGGGGGCCTGGCCATCTCCCTGGACAAAAGTGGGGTGCCTTTGCTACAGgta
gtggcacgggcctatagtttcagcatttgggaggtagaggcaggagaatcaggagttcaagcttatccttggcaacacacctagttt
aaggtcagcctgggctacatgagagcctacctcccccatcccctacCCCCAGAAAAGAAGGAAAATCTGGGGG
CACTGTGGATTTCTCCTCTCTTTTCTCTACCTGTTGAAAGCAAAGTCTAGGAAGGCCCCA
AACATGATAGCATTTGGGCCCTTAGTAAGCTGAAGATAAAAAGGAGAAGCTGTTTGGCTT
CGCCCCACAAAGCAGCTGCAGG

**IRAK4-mScarlet-i HDR template**

CTGCCGATCTCCCGGTCCCTCTTTTCGGTTCATCAGGTCTGAGTCCCTGTAGATAAAGAC

AAGGTGCCTGGATCCACAAGGTGGAATTCCAATATCAGACACACATCAAGAAGCCGATG
ACCCTGCTGGCAAACGCTGCCAGCAAGAGTGGGAGCAGCTAGCTCACCCCAGAAAGGC
AAGAATTATAAAGCACCTGTGCAGGCGCATATATGAATAAAGCCAGCAAATCTGTAAATG
TGTGCCTATGTATGCATTTCCTTTCGTAACATCTGACCTGTACAGCTCCTGATGAGTCTG
CCCCCCCAAAAAATACTGAGATAAATCGAAATAATTTACATTTATCTTCCGATTTCTTTAAC
TTGAAAGGGTTTAACCAAGTTTTAATATCGTTTTCAAGCTGGATATTAaagaagagattgaagatg
aagagaagacgattgaagattacacggatgagaagatgagCGATGCGGACCCTGCTTCGGTGGAAGCAA
TGTACTCTGCTGCTAGCCAGTGTCTGCATGAGAAGAAAAACAGACGGCCAGACATTGCA
AAGGTATTCACTTCCGAGTCCATCTACACAGCGAGGGGTGTGGTGCCTTATCCTTTACTA
AGTATCCATTTCAAGACATCGGCTTAAtCTaGTTCTTTTGATCTCTTcCTTcTTcAAAGGTTC
AACAGCTGCTACAAGAGATGTCTGCTGGAGGAAGTGGAGGTTCTGGTGGTAGTGTGAGC
AAGGGCGAGGCAGTGATCAAGGAGTTCATGCGGTTCAAGGTGCACATGGAGGGCTCCA
TGAACGGCCACGAGTTCGAGATCGAGGGCGAGGGCGAGGGCCGCCCCTACGAGGGCA
CCCAGACCGCCAAGCTGAAGGTGACCAAGGGTGGCCCCCTGCCCTTCTCCTGGGACAT
CCTGTCCCCTCAGTTCATGTACGGCTCCAGGGCCTTCATCAAGCACCCCGCCGACATCC
CCGACTACTATAAGCAGTCCTTCCCCGAGGGCTTCAAGTGGGAGCGCGTGATGAACTTC
GAGGACGGCGGCGCCGTGACCGTGACCCAGGACACCTCCCTGGAGGACGGCACCCTG
ATCTACAAGGTGAAGCTCCGCGGCACCAACTTCCCTCCTGACGGCCCCGTAATGCAGAA
GAAGACAATGGGCTGGGAAGCGTCCACCGAGCGGTTGTACCCCGAGGACGGCGTGCTG
AAGGGCGACATTAAGATGGCCCTGCGCCTGAAGGACGGCGGCCGCTACCTGGCGGACT
TCAAGACCACCTACAAGGCCAAGAAGCCCGTGCAGATGCCCGGCGCCTACAACGTCGA
CCGCAAGTTGGACATCACCTCCCACAACGAGGACTACACCGTGGTGGAACAGTACGAAC
GCTCCGAGGGCCGCCACTCCACCGGCGGCATGGACGAGCTGTACAAGTAAAACTAATA
AACttttttttttttttttttACTTCTCGAGAACCTGGAGACCGGAAAAGCACTTTGACACTGAGCTGCG
TCACCTACCTGTTGTTTGCATTTGTTTTTGTTTTCCTAAATGCGCTTTATGTTTAACATGGT
AGCATAGTGCAATGTACCAATGTACTGGTTAGTTAGTTAATAGACGAGTTGGCACCGCAG
CCAAAGTGATAGTTGCATGGTAGCACTTGGCCCCGCCCACCTAGTATGGGGCTCATGTT
CCACAGCAACATCACCAAACTATAGTGAACTTAAGATTCAGGAAAACAGCACTGTGCTAT
ACAGAGACAATAGGATTATTTGGGGCGGGGGAGTgagacacggtgtctctgtgtagccctggatgtcctg
gaacttgctctgtagaccaggctggccttgaactcagcaatctgcctgtctctgcctactgcgtgttgggattaaaggtgtgtgccactc
ctgtaaagctttggaccagctcttagctgccttaagccttagtgctggcttttggtttttgataggaacttcatgctcctgtaaatagctcctc
acctaggttcctgtaaggaacctctctccactgctcctgtaaccctaactaaactcattggttcaacaatgtagacatgagtggaatct
attcttctgtctgttattgtggtcctattttgaatgaatagatgtttgcttgtcat

**IRAK1-mScarlet-i-2A-PuroR HDR template**

GTTATCTGTCATTCTGTTGGCTGTATGGGGCCTGTGGTATTAACCTGAACATAGGCAAGT
CTAGGCCAACCTGAAGCAGCAGAGTAGCTGTGTGGTTTGACCACAGGGTGTCAGATCTA
GGCTGCCTTGCCCAGCCAGGAACCCATGCTCTGCCCATCCTGAGAGCCTCTCTCTCCAC
ACAGGTATACAAGAGACTAGAAGGGCTTCAGGCAGGGCCTCCCTGGGAGCTAGAGGTT
GCCGGCCATGGCTCCCCTTCCCCACAGGAGAACTCCTACATGTCTACCACTGGCAGTGC
CCAGAGTGGGGATGAACCATGGCAGCCTCTAGTAGTGACCACAAGAGCCCCAGCCCAG
GCTGCCCAGCAACTCCAGAGAAGTCCCAACCAGCCAGTGGAAAGTGATGAGAGTGTTCC
CGGCCTCTCTGCTACCCTGCATTCCTGGCACTTGACTCCAGGTTCCCACCCAAGCCCAG
CGTCCTTCAGAGAGGCTAGCTGTACCCAGGGAGGCACTACCAGAGAATCAAGTGTGAG
GAGTAGCCCAGGCTTCCAGCCTACAACCATGGAAGGTAACTCGGAATAGGATCTCCCTT
CTTAACCCTTCTCCCCACCCCATGCCTAAAATCAAAAGTGTAGAATTCTAGGTTCTGTCTA

GGCAGCCTGGGAGCCTGGTCTGTGATCTGATCTCTCCCAACCTGGGCCCCAGAATATAC
TGGAGAATTCTGCCCTGCAGCCAGCATCACTAGCAGGGGGGAGCTGGGAAGACTCTGG
AGGCTAGAGTGTCACTCAAGATTGCAGGGACAagccgggcgtggtggcgcatacctttaatctcagcacttg
ggaggcagaggcaggtggatttctgagtttgaggccagcctggtctacatacagagtgagttccaggacagccagggctacaca
gagaaaccctgtctcgaaaaaaccaaaaaaaaaaaaaaGCAGGGACAGCTAAGAAGTTACTATCTCTG
CTCAGGCTCACCCACGGGCAGCTCATCCCTGCTGTCATCAGAACCACCACAGATCATCA
TCAACCCAGCCCGACAGAAGATGGTACAAAAGCTGGCTCTTTATGAAGAAGGGGTCTTG
GATAGCCTGCAACTGCTGTCATCAGGCTTTTTCCCAGGTACCTGTGGATAACTCTAGCTC
AGCTTCCCCAGAAGAGCCTCCCACCGCTGTCATACACACATACAAACATGCATACAAAGT
CATACTCTGCAGCTCCAGGGCATACAGGCATAATTCCTTATGTCAACCCTAAACAGCAGA
ACATTTGTTCTGGCTGCACAATTAAGGCTCCACTTACCAAGAATATAGGACCCAGCTTTT
GCTCAACATCTGTAGGGAGATGCCTGGCTCAATCCCTTACTTCCCACCAGTAGCACTCC
CACATGCTCCTAGAAAATGTGAAGTTCCAGCAGGAATTTTCTACTCCAGGTACTATATGA
AAAGCTGCTGTTTAGCCAAGCATGTTCTGGGCCTAGGAATTGCTGGGATACCTTAGCTG
GGGATAAGAGTGTACCTGTAAAGATCATCCAAACTAGCAggccggagagattgcccagtgaataagg
caatgggctgcttttgcagatttacttggctttagtttccagactccacatttggtagttcacagctacttgtaactccagtcccaggggat
ctgaatccctcttctggcctccgcagatacctttgtatatacatgtatatactttgtatatacatgGTGTATTTAAAACTCACG
TAGAAACACACATATAAAAGGAAAAATCTTATTTTCAAAAAGCAAACTAAAGGTATAAAGA
AAACCTTGGCTCCAAAGACTGCCCTGagctgggcagtggtaacgcatgcctttaatcccagcacttgggaggca
gaaacaggcaagtttggggccagcttggtctacagagcaagttccaggatagccagggcaacagacaaaaaaaccctgtgag
aaaaAGGTGGGGTggggtagggaattagctcagttgtagagcatttgcttagcaagcgaagtgccctgagttccagtctcC
TTCCCACATGCTCAGTTTGGAGTGGGAGGGGGCCCACAAAGTTCTGTGCTCATGGTTCA
TGTCAGGGTATTTGAAAGACAAGAAGTTGTCCCCAACTTATTAAGTGATATTTTCTTTTCC
ATAGGCTTGGATTTAGAACCTGAAAAGtctCAGGGACCTGAAGAAAGTGATGAATTtCAGAG
CGGAGGAAGTGGAGGTTCTGGTGGTAGTGTGAGCAAGGGCGAGGCAGTGATCAAGGAG
TTCATGCGGTTCAAGGTGCACATGGAGGGCTCCATGAACGGCCACGAGTTCGAGATCGA
GGGCGAGGGCGAGGGCCGCCCCTACGAGGGCACCCAGACCGCCAAGCTGAAGGTGAC
CAAGGGTGGCCCCCTGCCCTTCTCCTGGGACATCCTGTCCCCTCAGTTCATGTACGGCT
CCAGGGCCTTCATCAAGCACCCCGCCGACATCCCCGACTACTATAAGCAGTCCTTCCCC
GAGGGCTTCAAGTGGGAGCGCGTGATGAACTTCGAGGACGGCGGCGCCGTGACCGTG
ACCCAGGACACCTCCCTGGAGGACGGCACCCTGATCTACAAGGTGAAGCTCCGCGGCA
CCAACTTCCCTCCTGACGGCCCCGTAATGCAGAAGAAGACAATGGGCTGGGAAGCGTC
CACCGAGCGGTTGTACCCCGAGGACGGCGTGCTGAAGGGCGACATTAAGATGGCCCTG
CGCCTGAAGGACGGCGGCCGCTACCTGGCGGACTTCAAGACCACCTACAAGGCCAAGA
AGCCCGTGCAGATGCCCGGCGCCTACAACGTCGACCGCAAGTTGGACATCACCTCCCA
CAACGAGGACTACACCGTGGTGGAACAGTACGAACGCTCCGAGGGCCGCCACTCCACC
GGCGGCATGGACGAGCTGTACAAGggttccggtGCCACGAACTTCTCTCTGTTAAAGCAAGC
AGGAGACgtggaagaaaaccccggtcctatgaccgagtacaagcccacggtgcgcctcgccacccgcgacgacgtcccc
cgggccgtacgcaccctcgccgccgcgttcgccgactaccccgccacgcgccacaccgtcgacccggaccgccacatcgagc
gggtcaccgagctgcaagaactcttcctcacgcgcgtcgggctcgacatcggcaaggtgtgggtcgcggacgacggcgccgcg
gtggcggtctggaccacgccggagagcgtcgaagcgggggcggtgttcgccgagatcggcccgcgcatggccgagttgagcg
gttcccggctggccgcgcagcaacagatggaaggcctcctggcgccgcaccggcccaaggagcccgcgtggttcctggccacc
gtcggcgtctcgcccgaccaccagggcaagggtctgggcagcgccgtcgtgctccccggagtggaggcggccgagcgcgccg
gggtgcccgccttcctggagacctccgcgccccgcaacctccccttctacgagcggctcggcttcaccgtcaccgccgacgtcga
ggtgcccgaaggaccgcgcacctggtgcatgacccgcaagcccggtgccTGATTTGTTCACTCTGACAAATCC
CTCAGACTCAGAGGTCAAAGTTCTCATGCTTGGAAGTTCTCATAGTGTGCAAACTCCTCA
GTAGCCCAAAGGCATCAGGTGTGGGGGCTCACTCTGGCAAAGGAGAGAGGGCAGTGGA

CCCTGCTGTTCTCAGAATGTGAACACAGACAGGGCCAGCCAAAGAACTTGATAGAAGCT
GATAAGGCAGGCAATGTGAGGCCAGCATTTGTGAGCAAGGCAAGCAGAGTACCTAAGGA
ATTGGGTAGTACAAAACAGAAGACTggggctggtgagatggctcagtgggtatgagcacccgattgctcttccaa
aggtccggagttcaaatcccagcaaccacatggtggctcacaaccatctgtaacaagatctgacgccctcttctggagtgtctgaa
gacagctacagtgtacttatatataataaataaatctttaaaaaaaaaaaaCAGAAGACTGCTAGCTACACAGGC
CCAGAGGACACAAGTAGGTCTGTTTGTATGGTGGGAGCTAGCTCCCCTGAGAGCTAGAT
AATATATCTGCAGTCTGTGAGTCTATTCCTCAGAACTTCCCTAAAACACCAAGCGAAAGTA
GCTGTTGTGCTTGTAAATTGCaaaaaaaaaaaaaaaaaaaaaaaaaaaaGAGTGTCTTACAGTTCAG
TCCACTAGCTAGAAGTGAGCACTCTGAGATTCTCCCAAGAGTCTAATAAAACCCCCCTCC
CCATCTTGGGTGGATATATGCAAGCC

**IRAK4-KO-BlastR-Stop-HDRtemp**

TTCTAGATGCTGTCTCTGAAACTGTTGTAAAAGCgttgttgttgtttgtttgtttgttgtttcaagacagggtttct
ctgtgtagccctggctgtcctggaactcactctgtagaccagactggcctcgaactcacaactcaaagacccacctgcctccccagt
gctggcattaaagacatgcgccacaaccacccggctGACTTGTTGTAAAGTTGTAATAGGAATCTTTCATA
TTTCAtGTAGAAGATGgccaagcctttgtctcaagaagaatccaccctcattgaaagagcaacggctacaatcaacagc
atccccatctctgaagactacagcgtcgccagcgcagctctctctagcgacggccgcatcttcactggtgtcaatgtatatcattttact
gggggaccttgtgcagaactcgtggtgctgggcactgctgctgctgcggcagctggcaacctgacttgtatcgtcgcgatcggaaat
gagaacaggggcatcttgagcccctgcggacggtgccgacaggtgcttctcgatctgcatcctgggatcaaagccatagtgaagg
acagtgatggacagccgacggcagttgggattcgtgaattgctgccctctggttatgtgtgggagggcTGATGATGAcAAC
AAGCtGTTGACACaATCGACATACATACGCAACCTTAATGTGGGGATCCTTAGGAAGCTG
TCGGATTTTATTGATCCTCAAGAAGGGTGGAAGAAATTAGCAGTAGCTATCAAAAAGCCG
TCCGGCGACGACAGATACAATCAGTTCCATATAAGGTAACTCCTCCTCCCCATCACAAAA
AGGATGCGGGCGTGACTACCATCTAACGGGACAGTTTAGAAGGTGAACCTTTCAATCGT
GGTGGTGCTATTGGTAATATGTAAGTGTTTTGTATATAGAATTTTTGCTAAAATTGCCAGC
ATGTCTGAGCTAAAGGTGTGAGCCCGCAAAGGTGTTTCTGAGCTAGAATCCAGCTCCCC
GTCTTTGTTCTAGTTCAGTTTGTTGGGCCAAGAACAGTTTATGCTGCTTCCGGCTTTGCAT
GTCCAAACTGAAGAGACAGAAAGCTATTGTAAGAATCTAATGAGATTTTAGAGTCTACACT
TTGCAGTGTGTGAGTAAGGATGAGTGCCCATGCGTCCTTACTCACCCAGGATTCTTGGAT
CTAGGATTTTCTAACAGCTTCCTGTAGTCGCCTCTCTTACATCTCTCCTCGATTCAGCAGA
AAACACAAGTTCGGATTTTAATGTTGTACTTTGTTCTGCAGTAAAACGCAAC

**IRAK1-KO-BlastR-Stop-HDRtemp**

CAGTAGAGGAATAGGCAAGAGACACTCATTCATGTGGCTAAAGGTCACAACAGGTGGTA
TCACCAACAACTACTCTAGAAAGGGGTAGGAGAAGCTGCTAGACCCTACTATCCTTATGG
AGAGAAGTGCAAACTAAGATAAAAGTGTGACAGAAAAAGGAAAGGGACATTTAGAGCCG
AACACCATGTGTATGTTAAGAGACATAAATGTGTTAGTTGGAAAGCTAATTAGAATCCTGC
ACAGGCAGATTCTGTGATCTAACACTTGCTGGCCCAGCTGCTGGAAGGTTCCTGGGGGC
GTGGTGGGGGCATGGAGATCTAGCACACAAGCAGTAGCATGGTATACACTCTTCCAGAC
CATTCTCAGGGCATCCATCCTTATCACCAAATGACTACAGTCTGCCAGAGGGCCATTTTG
TTACCCATATTTAACCCTGACTGACCAACAATTTGGCTGGCTCTCAAACTGCTCCCCAAC
TAAAGCGAGGGTAGCCTCTCCGTGACTTTCACCAGGCTTGGCATAAACTGCCTTTGTAGA
ACATCGTGTAAAAACATCGTGCCCTTTGTCTCTAGAACTGGAAGTCATTCTTCATCCTGTT
CAGCCCTATTCAGATGTTTGTGTAGGACCACAGACCTCGGTTCTCCCATTAGGGACTCCC
TCCTACGTAGCAGAGCTCCTTCCCCGCGTCAAGGGCGGCGCGGGACGCCCCCGTGTGG

CGGAAGGCCGCGGAGGGCAAGATGGCGGCGGGTCGAGCGCCGCCTCTTCCGCCCGGC
CGGGTGTCGCACGAGGCCGGCCTGAAGGGGAAGTGAGTCAGTGTCCGCGGACCCGGC
CCGCTAAGGCCCGCGCCCGCCGAGGCCCCGAGAGGCCTcggcccgggggcggcggcggcggcca
tggccaagcctttgtctcaagaagaatccaccctcattgaaagagcaacggctacaatcaacagcatccccatctctgaagactac
agcgtcgccagcgcagctctctctagcgacggccgcatcttcactggtgtcaatgtatatcattttactggggaccttgtgcagaact
cgtggtgctgggcactgctgctgctgcggcagctggcaacctgacttgtatcgtcgcgatcggaaatgagaacaggggcatcttga
gcccctgcggacggtgccgacaggtgcttctcgatctgcatcctgggatcaaagccatagtgaaggacagtgatggacagccga
cggcagttgggattcgtgaattgctgccctctggttatgtgtgggagggcTGATGATGAaCTAAGTGACGTACGAG
GTGCCACCCTGGGTTATGTGCCGCTTCTACAAAGTGATGGACGCCCTGGAGCCCGCCG
ACTGGTGCCAGTTCGGTGGGTGACGGCAGGCCGGCGTGGGGTGGTGGACTCAGGCTT
CTGGAGCTGAAGCCTGATGTTCCCCATCCCGCAGCGGCCTTGATCGTGCGCGACCAGA
CAGAGCTGCGGCTGTGCGAGCGCTCCGAGCAGCGCACAGCCAGTGTTCTGTGGCCCTG
GATCAACCGCAACGCGCGCGTAGCTGACCTCGTTCACATCCTCACGCACCTGCAGCTGC
TGCGTGCGAGGGACATCATCACAGCCTGTGAGCGAGCATAGCCCCAGGGACCCTAGGG
ATGGGAACCCAAGAGGGGTGACGGGTACAGAGTTTCCTGAGACAGGGAACCATATGGG
AACTcccccccccccaccccTTCACTTAGAGATAGAGAGCTCTGAGTCACAACCCTCCTTTCCTT
CCTTGTTATCCCAGGGCACCCTCCTGCCCCCGTTGTGCCCCCAAGCACCGCTGCCCCAA
GGCCCAGCAGCATCTCTGCAGGCTCTGAGGCCGGGGACTGGAGCCCCCGGAAATTGCA
GTCCTCTGCCTCCACCTTCCTCTCCCCAGGTAAAAAGAACCCGGGTGTTGGGCTTGATA
TGCCTGAGGAAAAGGCCACCATGACTGCTGACATGGCTGTAGACTGCTGCTGGTCTCTG
CCTCTCACTGCCATCTTTATGAAGCTTTTCCAGGCTCCCAGACCCATTCTGAGTCAGAGC
TCCTCCAGGTTCCACTCCCTGTTTCCCTCGGGCCACCACTACCATCTTCAGCCCCTTCCT
CCACCAAGGTAGGTGTCTCTTAACCCCCAGGG

Supplementary Figure 7

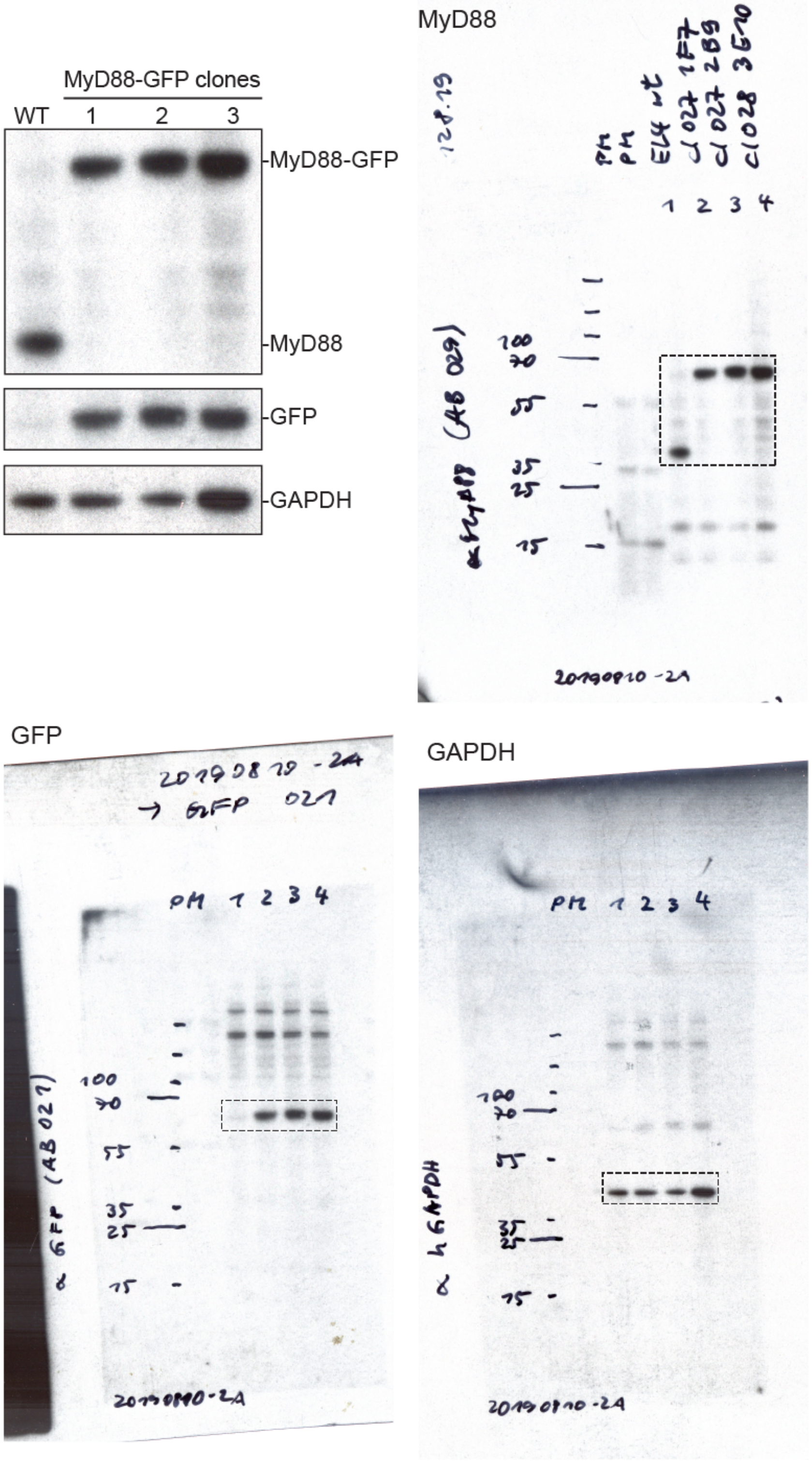


**Supplementary Figure 7. Full length Western blots from Fig. S1D.**

Supplementary Figure 8

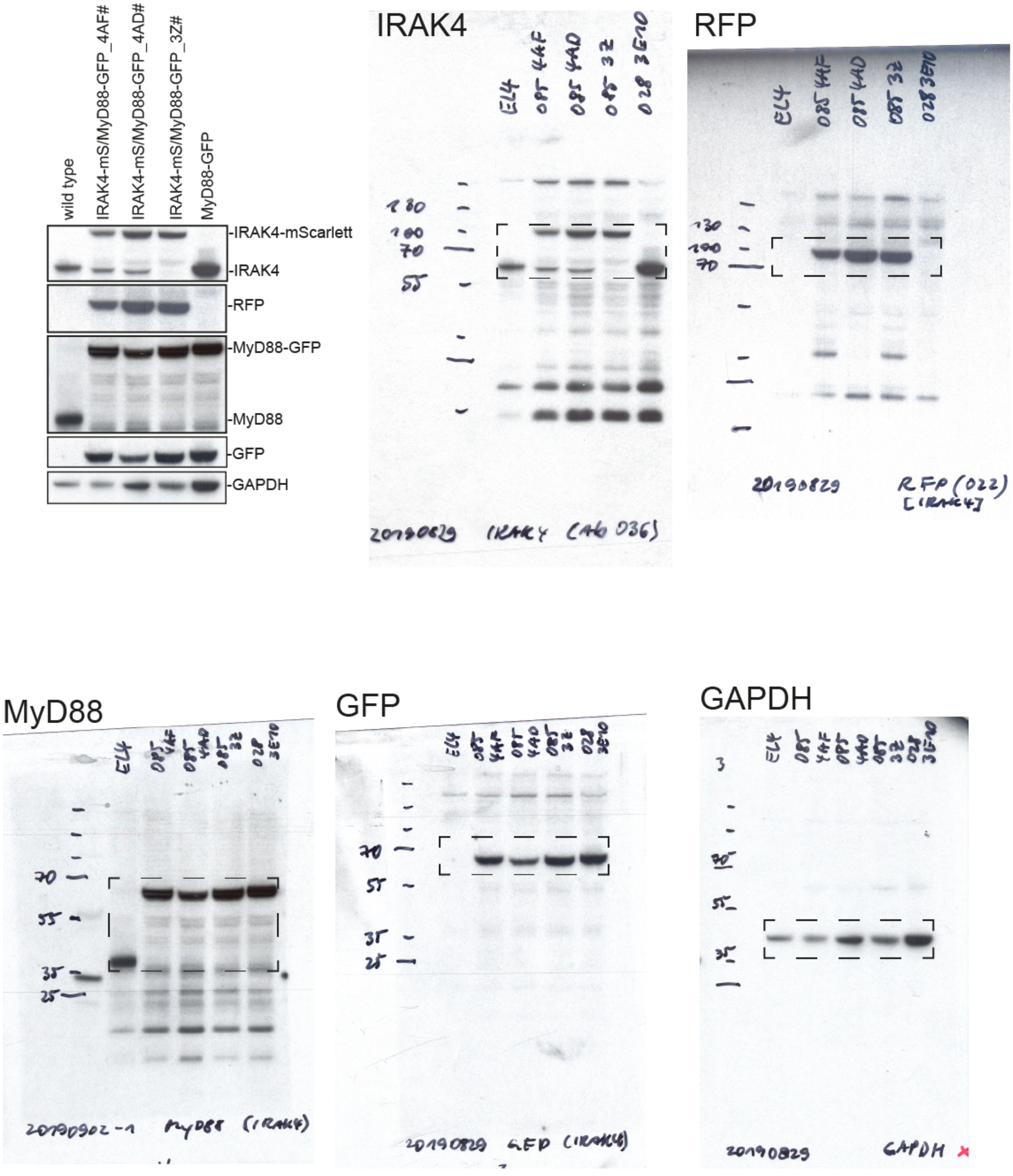


**Supplementary Figure 8. Full length Western blots from Fig. S4B.**

Supplementary Figure 9

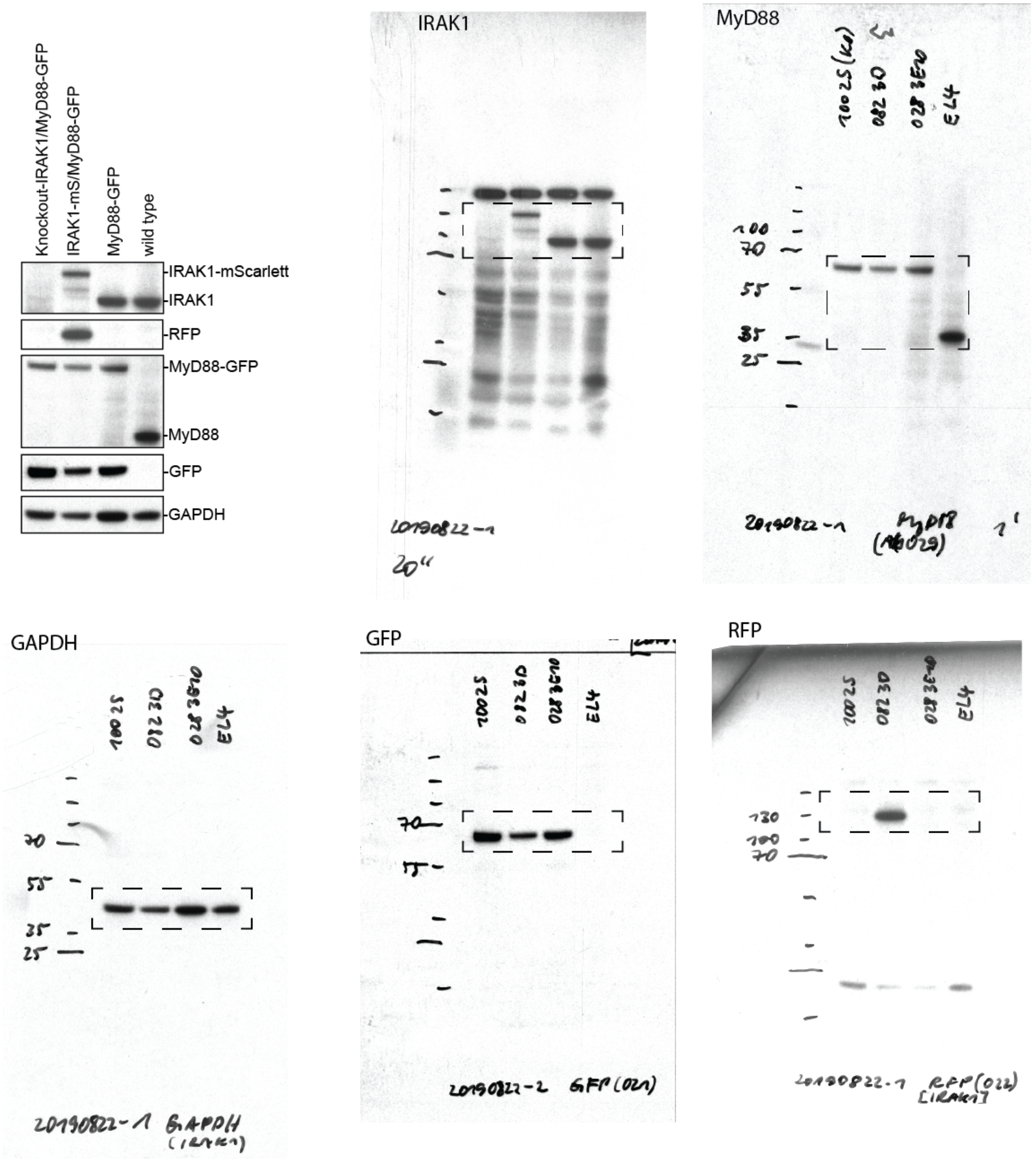


**Supplementary Figure 9. Full length Western blots from Fig. S4C.** Note WB images have been reflected around its vertical axis for Fig. S4C to enhance clarity.

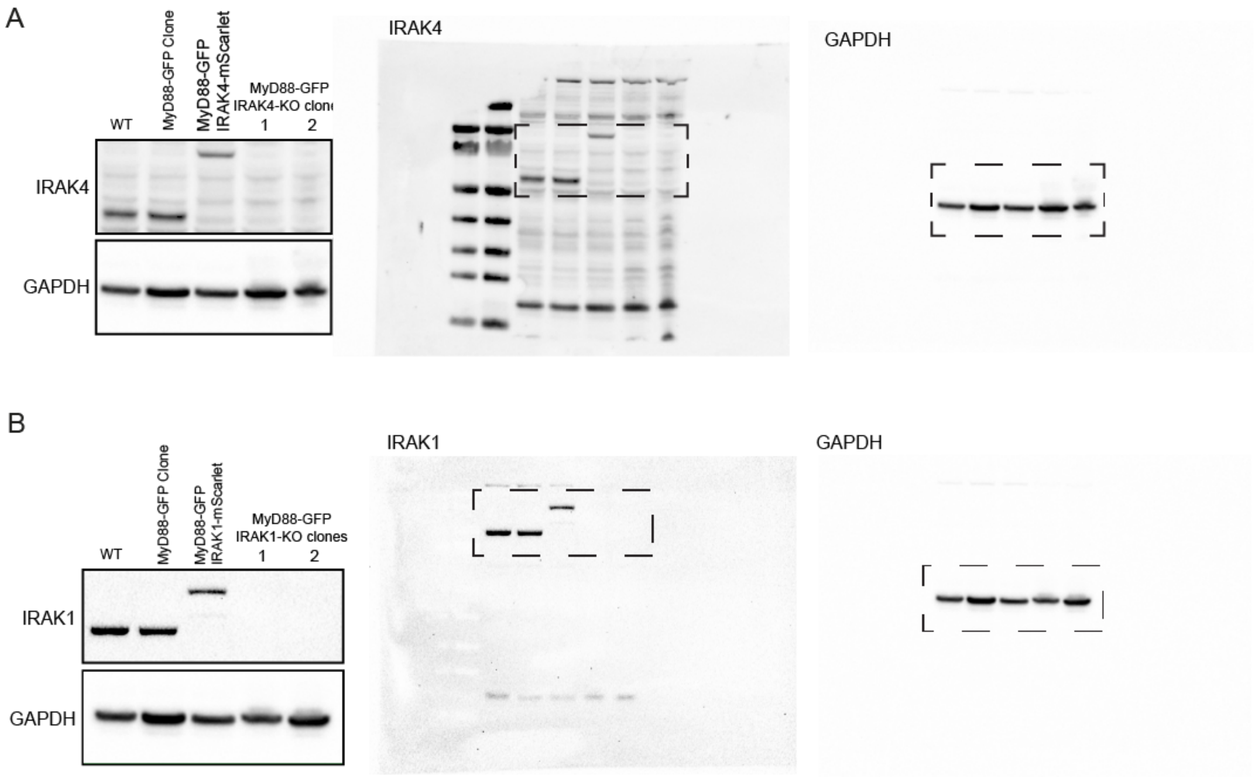


**Supplementary Figure 10. Full length Western blots from Fig. S6A.** Full length blot for Fig. S6A-B for IRAK4 and IRAK1 KO cells.
